# Heterochromatin-dependent transcription of satellite DNAs in the *Drosophila melanogaster* female germline

**DOI:** 10.1101/2020.08.26.268920

**Authors:** Xiaolu Wei, Danna G. Eickbush, Iain Speece, Amanda M. Larracuente

## Abstract

Large blocks of tandemly repeated DNAs—satellite DNAs (satDNAs)—play important roles in heterochromatin formation and chromosome segregation. We know little about how satDNAs are regulated, however their misregulation is associated with genomic instability and human diseases. We use the *Drosophila melanogaster* germline as a model to study the regulation of satDNA transcription and chromatin. Here we show that complex satDNAs (>100-bp repeat units) are transcribed into long noncoding RNAs and processed into piRNAs (PIWI interacting RNAs). This satDNA piRNA production depends on the Rhino-Deadlock-Cutoff complex and the transcription factor Moonshiner—a previously-described non-canonical pathway that licenses heterochromatin-dependent transcription of dual-strand piRNA clusters. We show that this pathway is important for establishing heterochromatin at satDNAs. Therefore, satDNAs are regulated by piRNAs originating from their own genomic loci. This novel mechanism of satDNA regulation provides insight into the role of piRNA pathways in heterochromatin formation and genome stability.

## INTRODUCTION

Repetitive DNA makes up a large fraction of eukaryotic genomes (Britten and Kohne 1968). Most repeat-dense genomic regions are gene-poor and tightly packed into heterochromatin (reviewed in Allshire and Madhani 2018; Janssen, et al. 2018). Tandem arrays of repeated sequences called satellite DNAs (satDNAs) are abundant in the heterochromatin of the pericentromeres, subtelomeres, and on sex chromosomes (Charlesworth, et al. 1994; Schmidt 1998; Schueler, et al. 2001). SatDNAs are typically viewed as selfish genetic elements that can spread rapidly in genomes and are generally repressed (Doolittle and Sapienza 1980; Orgel and Crick 1980). The de-repression of satDNA is associated with cellular senescence and various cancers (*e.g*. Ting, et al. 2011; Zhu, et al. 2011). However, satDNAs play roles in chromatin structure, chromosome segregation, and genome stability across wide range of taxa (Weiler and Wakimoto 1995; Dernburg, et al. 1996; Lippman, et al. 2004; Bouzinba-Segard, et al. 2006; Zhu, et al. 2011; Swanson, et al. 2013; Plohl, et al. 2014; Rosic, et al. 2014). SatDNA-derived transcripts have been detected in many species (Ugarkovic 2005; Usakin, et al. 2007; Biscotti, et al. 2015; Ferreira, et al. 2015). In insects, these transcripts may have roles in early embryos (Pathak, et al. 2013; Halbach, et al. 2020) and spermatogenesis (Mills, et al. 2019). Across organisms, satDNA-derived transcripts may generally be important for maintaining genome stability and integrity, yet the regulation and function of these transcripts remains poorly understood (reviewed in Janssen, et al. 2018).

Insights might come from the small RNA pathways that protect genome integrity by silencing repeats. These RNA interference pathways play roles in heterochromatin formation and maintenance at repeats across species (Hall, et al. 2002; Volpe, et al. 2002; Fukagawa, et al. 2004; Noma, et al. 2004; Verdel, et al. 2004; Novo, et al. 2020). In these pathways, small RNAs guide Argonaute proteins to cleave mRNA or silence genomic DNA by targeting complementary sequences (Hutvagner and Simard 2008). Among the most abundant types of repeat-derived small RNAs in animal germlines are the 23–32-nt PIWI-interacting RNAs (piRNAs) that target transposable elements (TEs)—genomic parasites that mobilize and can cause genome instability (Aravin, et al. 2006; Girard, et al. 2006; Grivna, et al. 2006; Lau, et al. 2006; Brennecke, et al. 2007; Houwing, et al. 2007). These piRNAs are particularly well-studied in Drosophila ovaries. The piRNA precursors are transcribed from discrete genomic loci containing primarily truncated TE sequences, called piRNA clusters. The piRNAs derived from these loci repress TE activity through both post-transcriptional (Brennecke, et al. 2007; Gunawardane, et al. 2007) and transcriptional silencing. In ovaries, piRNAs guide Piwi to genomic locations with complementary nascent RNAs and recruit heterochromatin factors to silence TEs (Wang and Elgin 2011; Sienski, et al. 2012; Le Thomas, et al. 2013; Rozhkov, et al. 2013).

There are two main types of piRNA sources in Drosophila ovaries—uni-strand and dual-strand piRNA clusters. Uni-strand piRNA clusters require promoter sequences and are either expressed only in somatic tissues (*e.g. flamenco*), or in both somatic tissues and the germline (*e.g. 20A*) (Brennecke, et al. 2007; Malone, et al. 2009; Mohn, et al. 2014). However, most piRNA clusters are heterochromatic dual-strand clusters, which are bidirectionally transcribed and do not necessarily require promoters (*e.g. 42AB, 80F*, and *38C1/2*; Brennecke, et al. 2007; Mohn, et al. 2014; Andersen, et al. 2017). Dual-strand piRNA clusters are expressed primarily in the germline, where their transcription is licensed by a non-canonical pathway that depends on the heterochromatin protein-1 (HP1) variant called Rhino (Rhi) (Klattenhoff, et al. 2009; Zhang, et al. 2014). Rhi recruits Deadlock (Del), an unstructured linker protein (Wehr, et al. 2006; Czech, et al. 2013), and Cutoff (Cuff), a protein related to the yeast Rai1 decapping enzyme (Pane, et al. 2011), to H3K9me3 chromatin. This complex is referred to as the Rhino, Deadlock, and Cutoff (RDC) protein complex (Mohn, et al. 2014). Moonshiner (Moon)—a paralog of the transcription factor TFIIA-L—interacts with Del and recruits TBP-related factor 2 (TRF2) to initiate transcription of dual-strand piRNA clusters (Andersen, et al. 2017). Most piRNA studies in Drosophila focus on their important role in repressing TE activity to protect genome integrity (*e.g*. Brennecke, et al. 2007). Given that TEs and satDNAs both are abundant repeats in heterochromatin whose activities are associated with genomic instability, we suspect that satDNAs may also be regulated by this piRNA pathway.

Consistent with our hypothesis, small RNAs derived from satDNAs exist in germlines (*e.g*. Aravin, et al. 2003; Saito, et al. 2006). However, little is known about these satDNA-derived small RNAs. Here we leverage publicly-available RNA-seq and ChIP-seq datasets and complement these data with cytological and molecular analyses of expression to study the regulation of satDNAs in the germline. SatDNAs are categorized based on their repeat unit size as simple (1-10bp) or complex (>100bp). We focus on two abundant families of complex satDNA in *D. melanogaster: Responder* (*Rsp*) and satellites in the *1.688* g/cm^3^ family (*1.688*). We show that complex satDNAs are expressed and processed primarily into piRNAs in both testes and ovaries. In ovaries, this expression depends on the RDC complex and Moon. Disruptions of the piRNA pathway leads to a loss of both satDNA-derived piRNAs and heterochromatin marks at satDNA loci. Our analyses suggest a model where the establishment of heterochromatin at satDNA is regulated by piRNAs originating from their own genomic loci. These findings add insight into the role of piRNA pathways in heterochromatin formation and genome stability.

## RESULTS AND DISCUSSION

### SatDNA transcripts originate primarily from genomic satDNA locus

To study satDNA expression patterns, we characterized transcripts from two representative complex satDNA families in *D. melanogaster—Rsp* and *1.688*—across tissues and developmental time points. *Rsp* consists of a dimer of two closely-related ~120-bp repeats in the pericentric heterochromatin on chromosome *2R* of *D. melanogaster* (Wu, et al. 1988; Pimpinelli and Dimitri 1989). The *1.688* family of repeats is the most abundant complex satDNA in *D. melanogaster* (Lohe and Roberts 1988). It comprises different subfamilies that exist as discrete tandem arrays in the pericentric heterochromatin named after their repeat unit sizes on chromosome *2L* (*260-bp*), chromosome *3L* (*353-bp* and *356-bp*), and the X chromosome (*359-bp*) (Losada and Villasante 1996; Abad, et al. 2000). Because there is high sequence similarity among these repeats, we analyzed all *1.688* subfamilies together unless stated otherwise.

We mined modENCODE datasets (Table S1 and Graveley, et al. 2011; Brown, et al. 2014) and found evidence for satDNA expression in total RNA-seq datasets from both sexes, and across different developmental stages (Figure 1; Figure 1–figure supplement 1). Both satDNA families are expressed in gonads, head, and other tissues (Figure 1A and Figure 1–figure supplement 1C). Their transcript abundance is low (RPM_*Rsp*_<10 and RPM_*1.688*_<300; Table S2), and generally increases throughout development and with adult age (Figure 1–figure supplement 1A and B). SatDNA-derived reads have very low abundance in the poly-A selected RNA-seq data (RPM_*Rsp*_<0.2 and RPM_*1.688*_<10; Table S2), indicating that the majority of satDNA transcripts are not polyadenylated.

**Figure 1.**
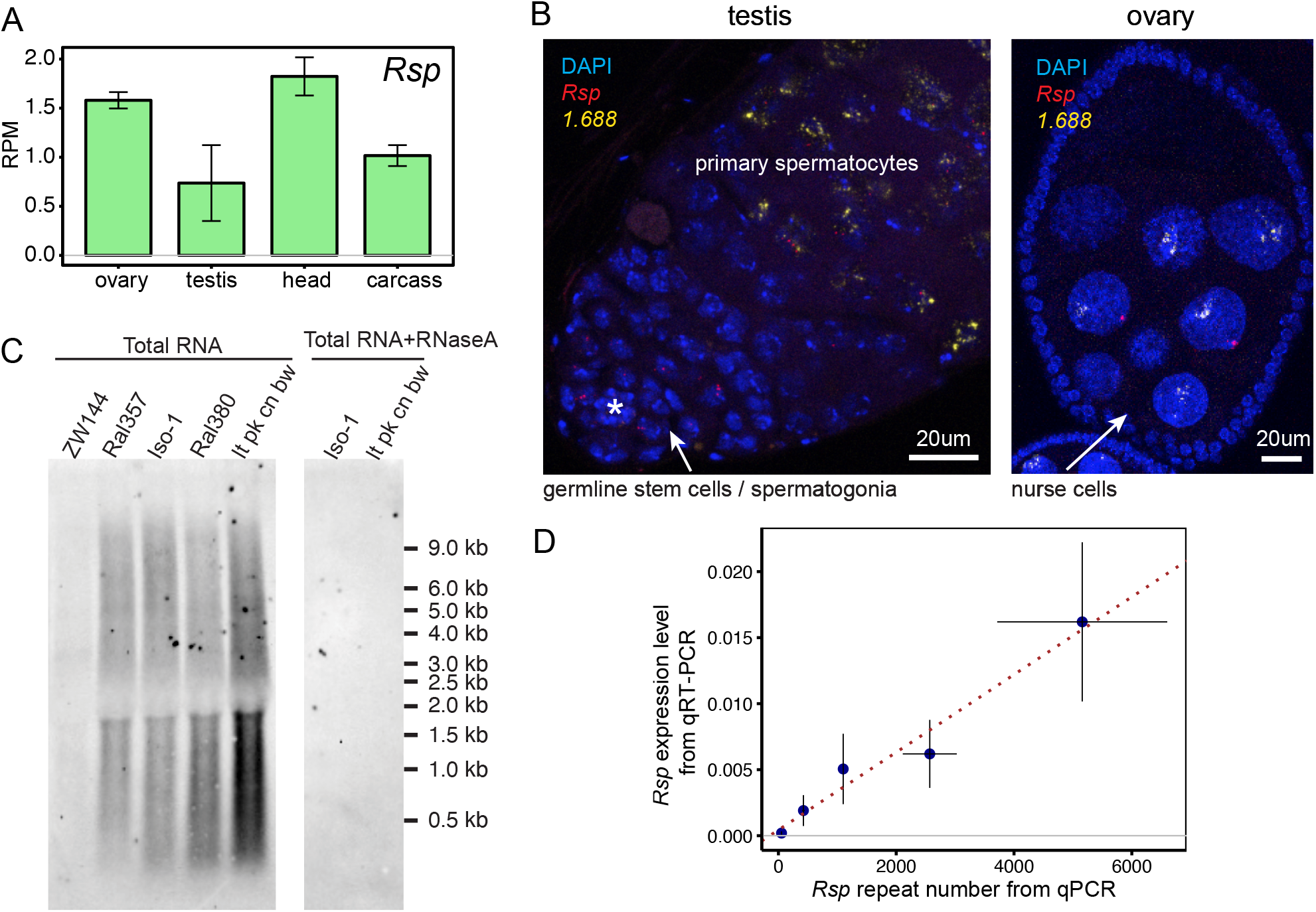
SatDNAs are expressed in ovaries and testes. (A) *Rsp* satDNA transcription level in various tissues (corresponding result for *1.688* is shown in Figure 1–figure supplement 1C). Carcass: whole body without the head, reproductive organs and digestive tract. Data from modENCODE (Graveley, et al. 2011; Brown, et al. 2014) (B) RNA FISH shows evidence for *Rsp* and *1.688*-derived transcripts in testes and ovaries, asterisk indicates the hub. The probe for *1.688* recognizes all of the subfamilies except for *260-bp* on chromosome 2L. (C) Northern blot probed with *Rsp*. Total RNA was extracted from ovaries of fly lines with varying copy numbers of *Rsp*: ZW144 (200 copies), Ral357 (600 copies), Iso-1 (1100 copies), Ral380 (2300), *lt pk cn bw* (4100). There is no signal after RNaseA treatment. Signal quantification (shown in Figure 1–figure supplement 1D) shows *Rsp* transcript abundance correlates with its genomic copy number (Pearson’s correlation coefficient r^2^=0.93, p-value=0.02); (D) qPCR and qRT-PCR quantification of *Rsp* copy number and expression level, respectively, of strains used in northern blot. A linear regression line is shown in the plot with red dotted line (Pearson’s correlation coefficient r^2^= 0.98, p-value=0.003). Details for (C) and (D) in Table S4. Figure 1–figure supplement 1. SatDNAs are expressed across tissues and developmental stages Figure 1–figure supplement 2. RNA FISH signals of satDNAs are from RNAs, not DNAs

To validate the presence of satDNA-derived transcripts in gonads, we used RNA fluorescence *in situ* hybridization (FISH). Both *Rsp* and *1.688* satellite transcripts are visible in testes and ovaries (Figure 1B; Figure 1–figure supplement 2A). These signals are undetectable after treating with RNaseA prior to probe hybridization (Figure 1–figure supplement 2B), which degrades single stranded RNAs, or RNaseH post probe hybridization (Figure 1–figure supplement 2C), which degrades the RNA in DNA-RNA hybrids. This suggests that these signals are from RNA rather than DNA. We detected satDNA transcript foci in ovarian nurse cells and in pre-meiotic testicular germ cells. Interestingly, in testes we detected *Rsp* signal at earlier stages of spermatogenesis (*i.e*., germline stem cells/spermatogonia) than the *1.688* signals (*i.e*. primary spermatocytes; Figure 1B). The difference in timing is notable, as *Rsp* is the specific target of *Segregation Distorter* (*SD*; Sandler, et al. 1959): a well-known male meiotic drive system that causes a defect in post-meiotic germ cells (reviewed in Larracuente and Presgraves 2012). *Rsp* transcription may therefore play some specific role in the male germline distinct from other complex satDNA.

The bulk of satDNAs are found in large blocks of tandem repeats in the heterochromatin with small blocks occurring in the euchromatin (Waring and Pollack 1987; DiBartolomeis, et al. 1992; Kuhn, et al. 2012; Sproul, et al. 2020). Some of the euchromatic (Menon, et al. 2014; Joshi and Meller 2017; Deshpande and Meller 2018) and heterochromatic loci in the *1.688* family (Usakin, et al. 2007) are transcribed. To determine if satDNA-derived transcripts originating from large heterochromatic loci is a general feature of other complex satDNAs, we examined transcript size and abundance in total RNA from ovaries of flies that vary in *Rsp* repeat copy number (Table S3; Khost, et al. 2017). We determined that, while transcript lengths were similar among these lines–ranging between <300 nt to > 9000 nt (Figure 1C)–the abundance of *Rsp* transcripts correlated with genomic copy number (Figure 1–figure supplement 1D and Table S4, Pearson’s correlation coefficient r^2^=0.93, p-value=0.02). We validated these hybridization results using qPCR and qRT-PCR to quantify *Rsp* genomic DNA and RNA transcript abundance, respectively (Figure 1D; Table S4, Pearson’s correlation coefficient r^2^=0.98, p-value=0.003). The correlation between genomic copy number and transcript abundance is consistent with most transcripts originating from the large blocks of heterochromatic satDNA.

### SatDNA transcripts are processed into piRNAs in Drosophila germline

Many different repeat-derived transcripts are processed into piRNAs (Aravin, et al. 2003; Saito, et al. 2006; Brennecke, et al. 2007) and endo-siRNAs (Czech, et al. 2008; Ghildiyal, et al. 2008; Okamura, et al. 2008; Menon, et al. 2014). To ask if complex satDNA-derived RNAs are processed into small RNAs, we re-analyzed published small RNA-seq data (Table S1; Ghildiyal, et al. 2010; Rozhkov, et al. 2010; Fagegaltier, et al. 2014; Mohn, et al. 2014; Quenerch’du, et al. 2016; Andersen, et al. 2017; Parhad, et al. 2017). We indeed detected satDNA-derived small RNAs in testes and ovaries (Figure 2–figure supplement 1 A and B). Our results suggest that the majority of these satDNA-derived small RNAs are piRNAs. First, these small RNAs are abundant in testes and ovaries, and their size distribution is typical for piRNA populations: an average of 90% of the RNAs range from 23nt to 28nt, with a peak at 24-26nt in *D. melanogaster* (Brennecke, et al. 2007; Figure 2A for *Rsp* and Figure 2–figure supplement 1C for *1.688*). Second, the satDNA-derived small RNAs bear a signature of the piRNA-guided RNA cleavage process called the ping-pong cycle. piRNAs amplified through ping-pong have a 10nt-overlap of antisense-sense piRNAs with a preference of uridine at the 5’ end (1U) or adenosine at nucleotide position 10 (10A) (Brennecke, et al. 2007; Gunawardane, et al. 2007). Our analysis of the ovary small RNA-seq data (Mohn, et al. 2014; Andersen, et al. 2017; Parhad, et al. 2017) confirms a ping-pong signature for satDNA-derived small RNAs: Z-score=4.55 for *Rsp* and 6.85 for *1.688* satellite (Figure 2–figure supplement 1E and G) and ~60-80% have 1U/10A (Figure 2–figure supplement 1D and F). Third, satDNA-derived small RNAs are bound by the PIWI proteins, as expected for piRNAs. Our re-analysis of published Piwi, Aubergine (Aub), and Argonaute3 (Ago3) RIP-seq data from ovaries (Brennecke, et al. 2007; Mohn, et al. 2015; Sato, et al. 2015) shows that *Rsp* and *1.688* RNAs interact with each of these proteins (Table S5). For example, ~0.9% and 0.1% of Piwi bound RNAs map to *1.688* and *Rsp*, respectively. For comparison, ~2% and 17% Piwi bound RNAs mapped to the dual-strand piRNA clusters *80F* and *42AB*, respectively. In contrast, only an average of 0.0005% of the reads from Piwi RIP-seq data mapped to miRNAs, which are abundant small RNAs not known to be bound by Piwi. This suggests that the abundance of satellite RNA in the RIP-seq data is not likely due to noise or contamination. Our results from Aub and Ago3 RIP data are similar to Piwi (Table S5; *e.g*. 3.1% and 0.1% of Aub-bound RNAs map to *1.688* and *Rsp*, respectively; and 1.8% and 0.07% of Ago3-bound RNAs map to *1.688* and *Rsp*, respectively). Together, these results indicate that satDNA-derived transcripts are processed into piRNAs in the female germline.

**Figure 2.**
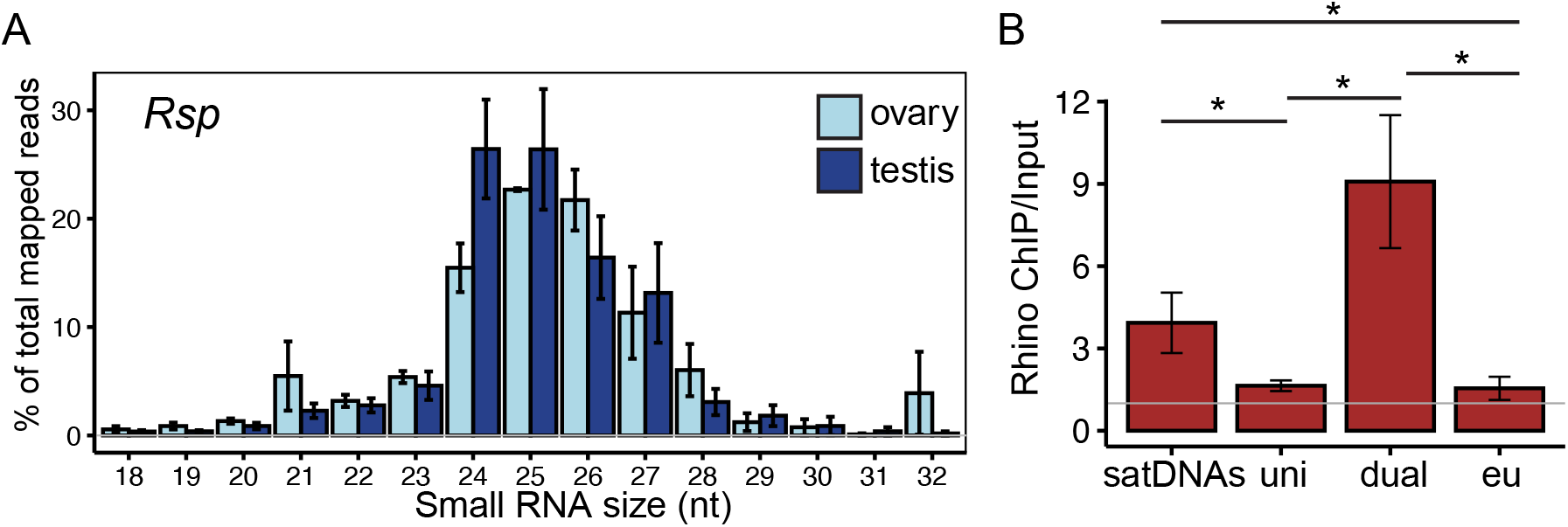
SatDNA produce small RNAs in *D. melanogaster* ovaries. (A) Size distribution of *Rsp* small RNAs in testes and ovaries (*1.688* distribution is in Figure 2–figure supplement 1C). (B) Rhino ChIP-seq result from ovaries showing the enrichment scores for satDNAs, uni-strand (uni) piRNA clusters, dual-strand (dual) and euchromatin (eu). The enrichment scores for each satDNA and piRNA cluster are shown in Figure 2– figure supplement 5A. P-values are estimated by pairwise t-tests with FDR correction (Benjamini 1995). * adjusted p-value<0.05. Figure 2–figure supplement 1. Satellite derived RNAs are mainly processed into piRNAs in the germline Figure 2–figure supplement 2. Non-uniform distribution of piRNA reads along satDNA consensus sequences Figure 2–figure supplement 3. Non-uniform distribution of piRNA reads along germline-dominant TE consensus sequences Figure 2–figure supplement 4. SatDNAs are transcribed from both strands Figure 2–figure supplement 5. ChIP-seq result shows the chromatin state of satDNAs, uni-strand piRNA clusters (*20A* and *flamenco*) and dual-strand piRNA clusters (*42AB, 80F* and *38C1/2*) and euchromatin

We examined the piRNA distribution along individual repeat units for *Rsp* and two subfamilies of *1.688 (359-bp* and *260-bp*) by blasting the corresponding sequencing reads to each consensus sequence. We find that the distribution of piRNA read depth is not uniform along the length of single monomers (*359-bp* and *260-bp*) or dimers (*Rsp*) of these satDNA repeats (Figure 2–figure supplement 2A and B). This pattern could arise if these piRNAs derive from repeat fragments overrepresented in the genome. However, when we look at the alignment depth of all genomic repeat variants, it is more uniform across the monomers/dimers than the piRNA pileup (Figure 2–figure supplement 2C). We observe similar non-uniform patterns of piRNA pileup over germline-dominant TEs (*e.g. invader6, mdg3* and *Het-A*; Figure 2–figure supplement 3), suggesting that these uneven distributions may arise from piRNA processing. The piRNA read pileup pattern also differs between ovaries and testes (Figure 2–figure supplement 2A and B), suggesting that there may be differences in transcription machinery, precursor production, or precursor processing between these tissues.

### SatDNA transcription resembles dual-strand piRNA clusters

*D. melanogaster* ovarian piRNAs originate primarily from uni- or dual-strand piRNA clusters. To determine which pathway controls the expression of satDNA-derived piRNA precursors, we asked whether transcripts come from one or both strands. We mapped total RNAseq reads from ovary and testis to the genome assembly. Collectively, for all genomic copies of *Rsp* or *1.688* satDNA (all subfamilies), we find a nearly 1:1 ratio of reads mapping to the plus and minus strands (Figure 2–figure supplement 4A; all mapped and uniquely mapped reads). However, the highly repetitive nature of satDNAs makes confidently assigning satellite-derived reads to a genomic location difficult. We therefore take advantage of our assemblies for two representative satDNA loci: the major *Rsp* locus on chromosome 2R and the *260-bp* locus, a subfamily of *1.688*, on chromosome 2L (Khost, et al. 2017). For these two loci, we confirm that reads map uniquely to both strands of the contigs (Figure 2–figure supplement 4B). Together, these results suggest that satDNAs are transcribed from both strands, similar to dual-strand piRNA clusters.

Dual-strand piRNA clusters are associated with the heterochromatin binding protein Rhi (Klattenhoff, et al. 2009; Zhang, et al. 2014). We therefore re-analyzed publicly available ChIP-seq datasets from ovaries (Mohn, et al. 2014; Zhang, et al. 2014; Parhad, et al. 2017) to determine if satDNA regions are also Rhi-associated. Our results for piRNA clusters are consistent with previous studies (Klattenhoff, et al. 2009; Mohn, et al. 2014; Andersen, et al. 2017): the dual-strand piRNA clusters have higher Rhi enrichment (mean enrichment ChIP/Input E_dual_=9.08) compared to uni-strand piRNA clusters (E_uni_=1.69; pairwise t-test with Benjamini Hochberg; Benjamini and Hochberg 1995) adjusted p-value P_adj_=0.01) and euchromatic genes (E_euch_=1.55; P_adj_=0.01). We found that complex satDNAs are in the top 30% of all repeats enriched in Rhi (full data in Supplementary File 1). The level of Rhi enrichment for satDNAs (E_sat_=4.70) is intermediate between the highly enriched dual-strand piRNA clusters (P_adj_=0.1) and the minimally Rhi enriched uni-strand piRNA clusters (P_adj_=0.01) or euchromatin (P_adj_=0.01 Figure 2B and Figure 2–figure supplement 5A). Unlike the uneven distribution of piRNAs along satellite monomers/dimers (Figure 2–figure supplement 2A and B), the distribution of Rhi ChIP-seq reads (Figure 2–figure supplement 2D) is similar to the alignment depth of genomic repeats (Figure 2–figure supplement 2C). This suggests that Rhi localizes to the large satDNA genomic loci, rather than a subset of smaller clusters or repeats across the genome (*e.g*. the 12 copies of *Rsp* inside an intron of *Ago3* on chromosome 3L; Figure 2–figure supplement 2C) or in potentially unannotated piRNA clusters.

### SatDNA transcription is regulated by RDC complex and Moon

Because we find that satDNAs generate piRNAs in the female germline and their chromatin is associated with Rhi, we asked if the same transcription and RNA processing machinery are used by both satDNAs and dual-strand piRNA clusters. We used publicly available small RNA-seq datasets generated from mutants of genes involved in the heterochromatin-dependent transcription initiation of dual-strand piRNA clusters: Rhi, Cuff, Del (RDC), and Moon (Klattenhoff, et al. 2009; Pane, et al. 2011; Czech, et al. 2013; Le Thomas, et al. 2014; Mohn, et al. 2014; Andersen, et al. 2017; Parhad, et al. 2017). We normalized piRNA abundance to the number of reads mapped to either miRNAs (Figure 3A) or the uni-strand *flamenco* cluster (Figure 3–figure supplement 1), neither of which should be affected by mutations in the RDC pathway.

**Figure 3.**
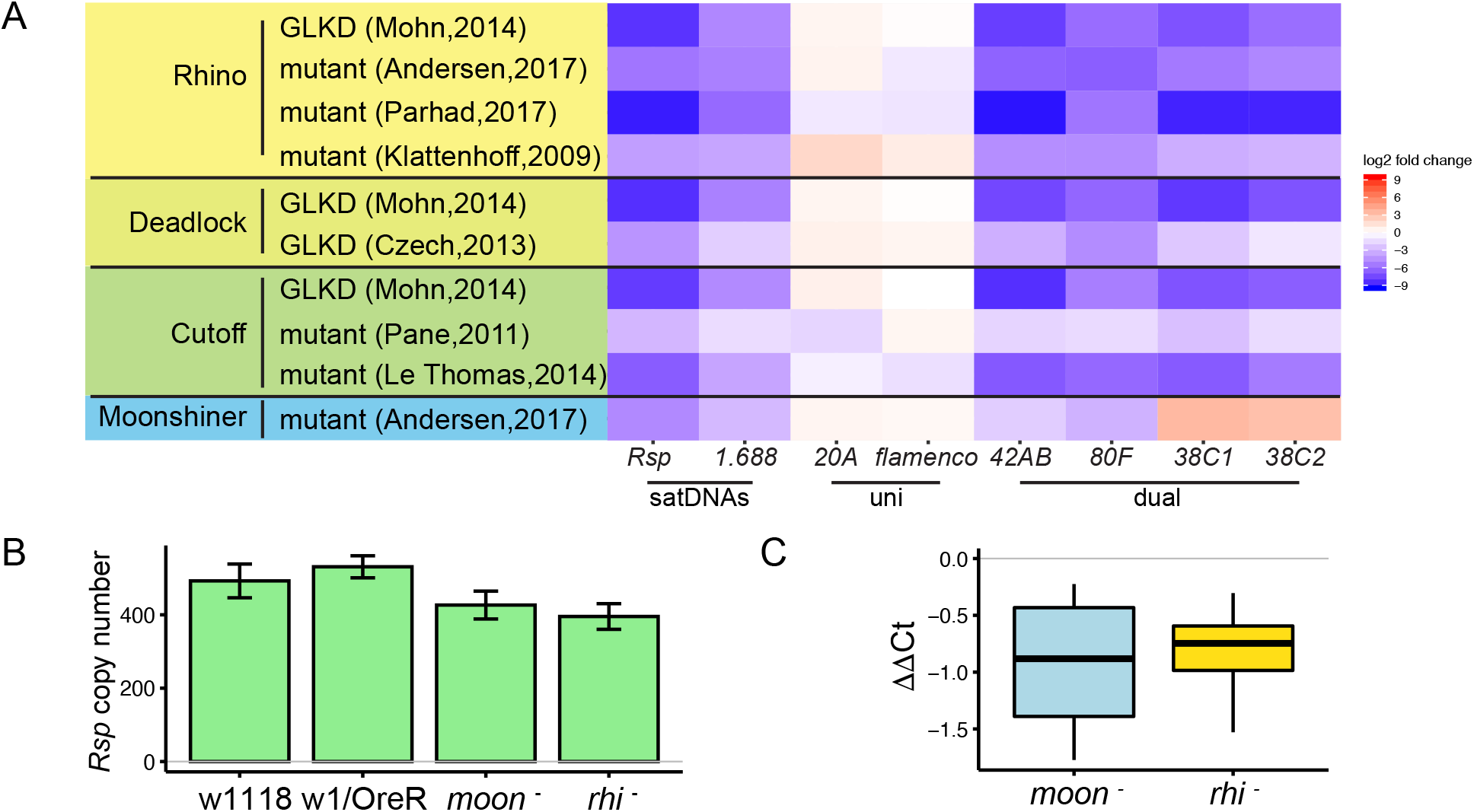
SatDNA loci are regulated by the heterochromatin-dependent transcription machinery in Drosophila ovaries. (A) Heatmap showing the quantification of piRNA abundance in small RNA-seq data from mutants of *rhino, cutoff, deadlock*, and *moonshiner* for satDNAs and piRNA clusters, normalized by miRNA level. GLKD: germline knockdown. Complete list of log2 fold changes in Table S6. (B) qPCR estimate of *Rsp* copy number in wild types and mutants. (C) qRT-PCR estimate of *Rsp* transcript level in mutants compared to wild types. ΔΔCt = ΔCt(wild type) - ΔCt(mutant), a negative value indicates lower expression in mutant. Student’s t-test, p-value=0.077, 0.048. Figure 3-figure supplement 1. SatDNA loci are regulated by the heterochromatin-dependent transcription machinery in Drosophila ovaries Figure 3-figure supplement 2. RDC and Moon mutants affect piRNA precursor transcription at piRNA clusters Figure 3-figure supplement 3. SatDNA piRNA production is affected in mutants of pathways involving piRNA precursor export, primary piRNA biogenesis, and the ping-pong cycle

Our analysis of the known piRNA clusters agrees with published results: the dual-strand piRNA clusters *42AB* and *80F* are Rhi- and Moon-dependent, and *38C1/2* is Rhi-dependent but not Moon-dependent. The uni-strand piRNA clusters *20A* and *flamenco* are not dependent on either protein (Klattenhoff, et al. 2009; Pane, et al. 2011; Mohn, et al. 2014; Andersen, et al. 2017). We find that the pools of complex satDNA-derived piRNAs are also reduced in RDC and Moon mutants (Figure 3A; Figure 3–figure supplement 1). In *rhi* mutants, *Rsp* piRNA abundance is 0.2–6.3% their levels in wild type datasets. Similarly, piRNA abundance for *1.688* is 1.4-7.8% their levels in wild type datasets (complete list of log2 fold change for satDNAs in Tables S6 and S7). The reduction in satDNA piRNA level is robust to normalization method (miRNA in Figure 3A; *flamenco* cluster in Figure 3–figure supplement 1). While the expression of simple satellite repeats like AAGAG was not decreased in these mutants (Table S6 and Table S7), the low abundance of AAGAG reads (the number of reads mapping to AAGAG are only ~0.5% of *Rsp* and ~0.03% of *1.688*) and known sources of bias for simple repeats (*e.g*., PCR bias in RNA-seq library preparation; Wei, et al. 2018) points to the need for different approaches to verify this finding. Overall, our results indicate that piRNA production from complex satDNAs is regulated by the heterochromatin-dependent transcription machinery associated with dual-strand piRNA clusters.

To further examine how the RDC complex and Moon affect complex satDNA transcription, we re-analyzed total RNA-seq data of the corresponding mutants (Mohn, et al. 2014; Andersen, et al. 2017). RDC and Moon mutants affect piRNA precursor transcription at the dual-strand piRNA clusters *42AB* and *80F* (Mohn, et al. 2014; Andersen, et al. 2017). Consistent with published reports, we detected decreases in steady state long RNA transcript levels at dual-strand piRNA clusters (Figure 3–figure supplement 2). However, we did not observe a significant decrease in steady state long RNA transcript levels for satDNAs (Table S8). To confirm this finding, we performed qRT-PCR using total RNA from ovaries of *rhino* (*rhi-*) and *moonshiner* (*moon^-^*) mutants (Andersen, et al. 2017). After controlling for genomic repeat copy number with qPCR (Figure 3B), *Rsp* expression level is lower, but not significantly so in *rhi* and *moon* mutants compared to wildtype (p-value=0.048 and 0.077; Figure 3C). Because satDNAs have generally low expression levels (*Rsp* and *1.688* total RNA levels are ~3% and ~25%, respectively, of both *42AB* and *80F*), we may have insufficient power to detect decreased expression in the mutants. It is also possible that the signal is masked by non-precursor transcripts. That is, there may be two kinds of transcription at satDNA loci: *1*) RDC-regulated transcription that generates non-polyadenylated piRNA precursors and, *2*) non-precursor transcription which is not well characterized and may also largely lack polyadenylation. In this context, it would be difficult to distinguish precursor from non-precursor transcripts derived from satDNA. However, when we re-analyzed the total and poly-A selected RNA-seq data from the Rhi mutant (ElMaghraby, et al. 2019), we find that the abundance of poly-A transcripts (which are likely a subset of non-precursors) is increased for *Rsp* and unchanged for *1.688* (Table S9) relative to wild type. This result suggests that changes in piRNA precursor levels may be masked by the non-precursor levels, similar to reports on piRNA cluster transcription in embryonic *piwi* knockdown ovaries (Akkouche, et al. 2017). This situation might arise if only a subset of satDNA repeats are RDC-regulated. Alternatively, the proportion of piRNA precursor-to-non-precursor transcripts in these mutants might shift such that the abundance of piRNA precursors decreases but the total RNA level does not.

We also asked if the satDNA-derived piRNA pool is affected in mutants of twelve genes involved in piRNA precursor export from the nucleus, primary piRNA biogenesis, and the ping-pong cycle (Figure 3–figure supplement 3; Czech, et al. 2018; datasets from Malone, et al. 2009; Handler, et al. 2011; Olivieri, et al. 2012; Preall, et al. 2012; Zhang, et al. 2012; Czech, et al. 2013; Sato, et al. 2015; Wang, et al. 2015; Table S1). For each of the datasets analyzed, we recapitulate previously reported results for all known piRNA clusters (Figure 3–figure supplement 3; Czech, et al. 2018). Our re-analysis of these data suggests that piRNA production for all complex satDNA is regulated by the primary piRNA pathway (Gasz, Vreteno, Shutdown), UAP56, and the ping-pong pathway (Ago3, Krimper). Some of our reanalysis results varied between datasets from different studies for satDNAs. For example, satDNAs show decreased piRNA levels in one mutant Zucchini dataset (Olivieri, et al. 2012) but increased levels in an independent Zucchini dataset (Malone, et al. 2009; Handler, et al. 2011). While further work is required to determine all of the components involved in processing satDNA transcripts, our results suggest that piRNA production at satDNA loci is regulated by the dual-strand piRNA pathway.

### Heterochromatin establishment at satDNAs requires Piwi

Consistent with their Rhi enrichment, we find that satDNAs are enriched for H3K9me3 in ovaries (Figure 2–figure supplement 5B; datasets from Klenov, et al. 2014; Le Thomas, et al. 2014; Mohn, et al. 2014). Piwi plays an important role in establishing H3K9 methylation on euchromatic TEs in ovaries (Mohn, et al. 2014) and heterochromatin more generally in embryos (Akkouche, et al. 2017). Transiently knocking down *piwi* expression early in the embryonic germline leads to a general decrease in H3K9me3 in the adult ovary, and a specific decrease in piRNA production and increase in spliced non-precursor transcripts at dual-strand piRNA clusters (Akkouche, et al. 2017). We therefore re-analyzed H3K9me3 ChIP-seq data from embryonic *piwi* knock down ovaries (Akkouche, et al. 2017). We detected a decrease of H3K9me3 at satDNAs (Figure 4A), suggesting that Piwi is also required for the establishment of heterochromatin at these loci. Consistent with the decrease in H3K9me3, piRNA production from satDNAs is also reduced (with some variation among replicates observed for *Rsp*; Figure 4B); and satDNA total RNA levels are increased (Figure 4–figure supplement 1), similar to dual-strand piRNA clusters (Akkouche, et al. 2017). However, it is again difficult to distinguish between satDNA precursor and non-precursor RNAs.

**Figure 4.**
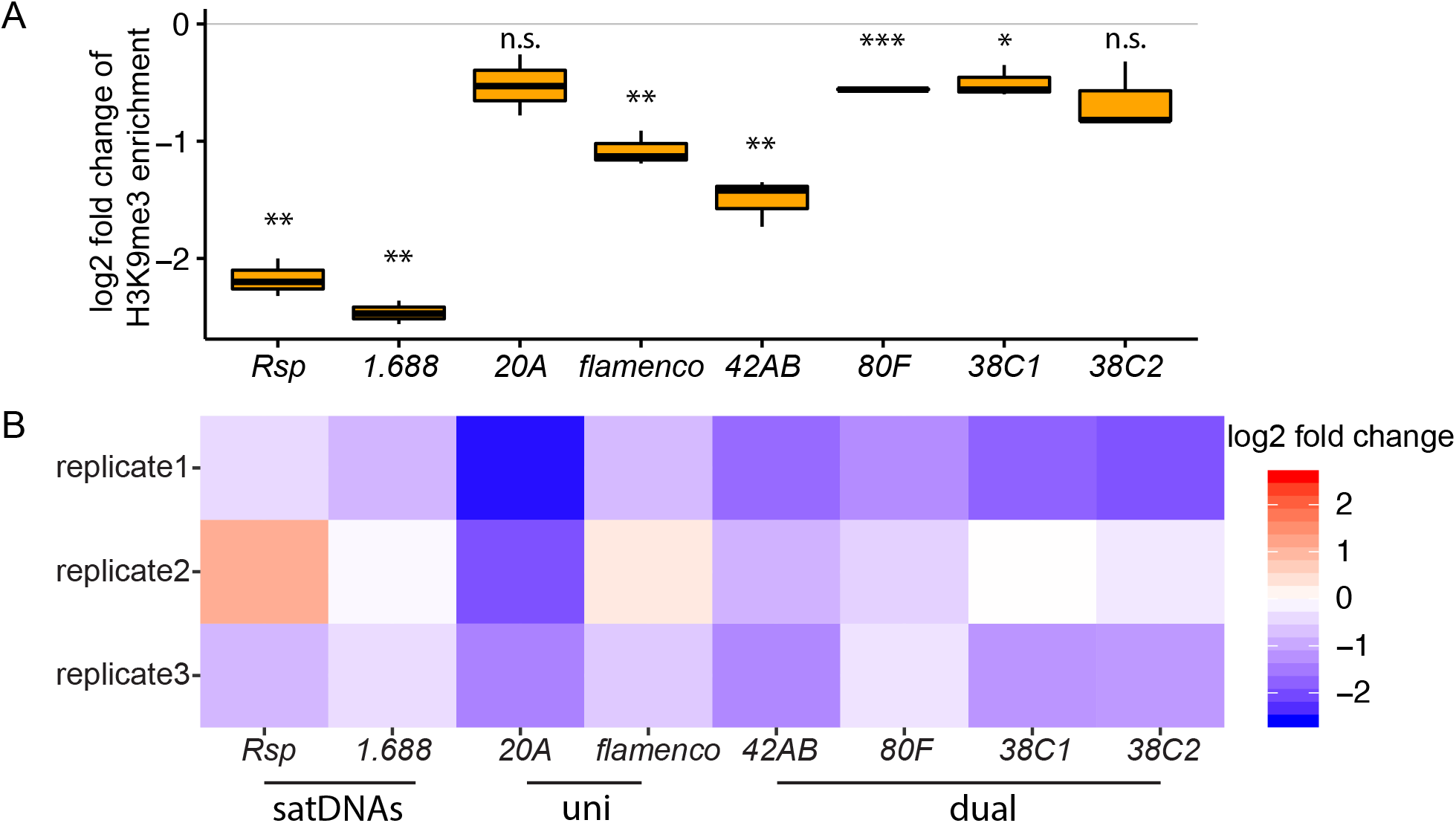
Heterochromatin establishment disrupted at satDNAs in *piwi* embryonic knockdown ovaries. (A) Log2 fold change of H3K9me3 ChIP/input enrichment shows satDNA H3K9me3 levels decrease in *piwi* embryonic knockdown ovaries compared to control. P-values are estimated by one sample t-test (mu=0) with FDR corrections (Benjamini and Hochberg 1995). * adjusted p-value<0.05, ** adjusted p-value<0.01, *** adjusted p-value<0.001. (B) Log2 fold change of small RNA abundance shows satDNA small RNA levels decrease, with variation observed for replicate2. Small RNA abundance is normalized to the number of reads mapped to miRNAs. Figure 4–figure supplement 1. Log2 fold change of total RNA abundance shows satDNA long RNA levels increase in *piwi* embryonic knockdown ovaries compared to control. Figure 4–Source Data. Log2 fold changes of small RNA abundance in *piwi* embryonic knockdown ovaries compared to control Figure 4–figure supplement 1-Source Data. Log2 fold changes of total RNA abundance in *piwi* embryonic knockdown ovaries compared to control

While Piwi is important for heterochromatin establishment at piRNA clusters, it appears to be dispensable for heterochromatin maintenance (Czech, et al. 2018). Depleting Piwi in the nucleus with *piwi* mutants lacking a nuclear localization signal (NLS; Klenov, et al. 2014), or knocking down germline *piwi* (Le Thomas, et al. 2013; Mohn, et al. 2014) affects H3K9me3 level on a subset of active transposons, but not on piRNA clusters (Klenov, et al. 2014; Mohn, et al. 2014). Similar to piRNA clusters, our re-analysis of these data show that the level of H3K9me3 on satDNAs is largely unchanged in the knockdown or mutant ovaries (with some variation observed among datasets; TableS10). These analyses suggest a role for Piwi in establishing, but not maintaining, heterochromatin at satDNAs in early embryos, which is important for producing piRNAs later in adult ovaries.

## CONCLUSIONS

piRNA pathways are primarily studied for their conserved role in protecting genome integrity by repressing TE activity in different organisms (Aravin, et al. 2006; Girard, et al. 2006; Grivna, et al. 2006; Lau, et al. 2006; Brennecke, et al. 2007; Houwing, et al. 2007; reviewed in Parhad and Theurkauf 2019). However, our findings support a more general role for these pathways. Here we show that transcription from satDNAs is regulated by the heterochromatin-dependent RDC machinery and Moon in ovaries and these transcripts are processed into piRNAs. Thus, complex satDNA transcription is regulated in a manner similar to dual-strand piRNA clusters in the female germline (Figure 5).

**Figure 5.**
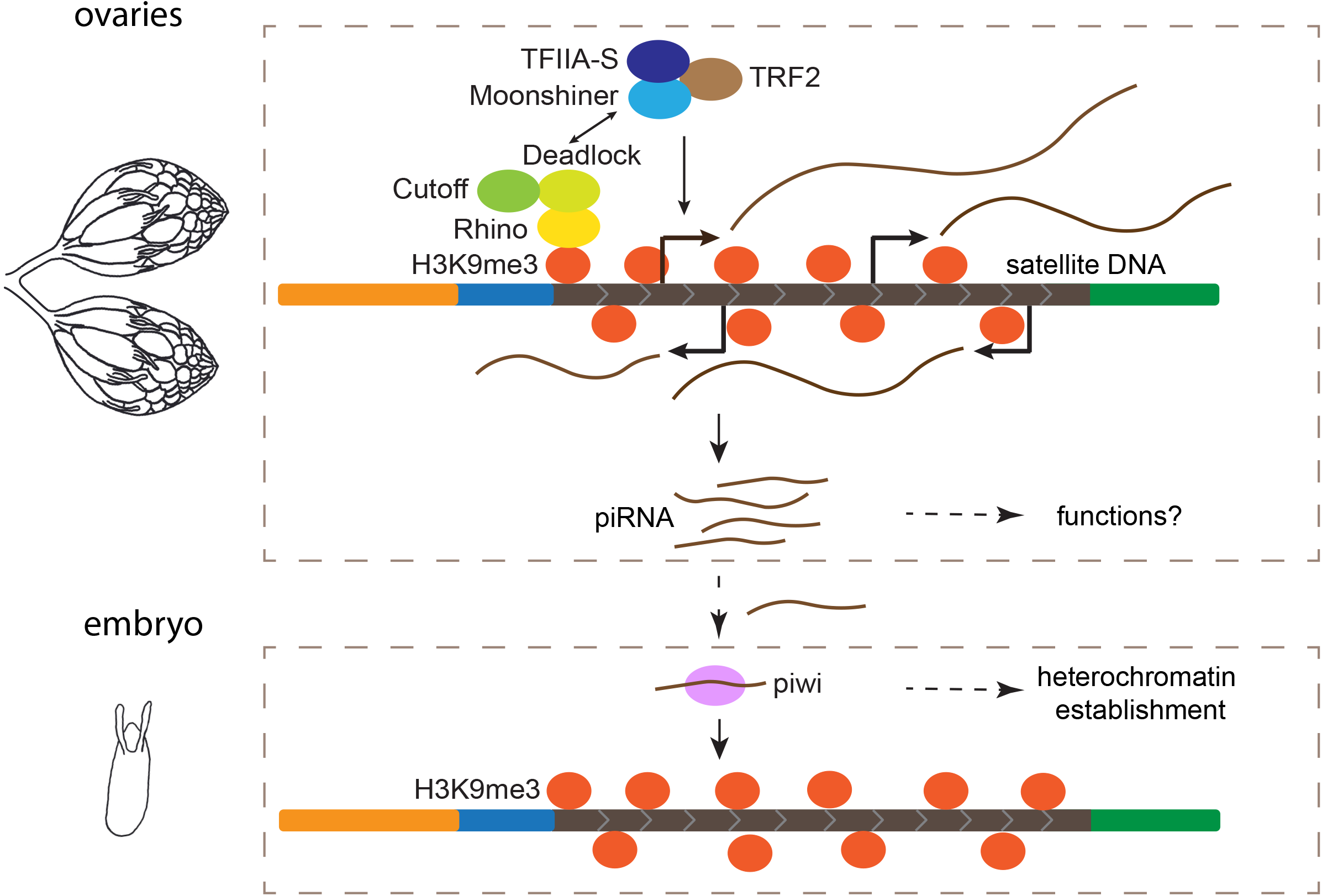
Model for maintenance of satDNA chromatin in female germline. Complex satDNA transcription is regulated by the heterochromatin-dependent Rhino-Deadlock-Cutoff and Moonshiner machinery, and the long RNA transcripts are processed into piRNAs. While their functions in ovaries are unclear, these piRNAs play roles in the establishment of heterochromatin at their own genomic loci in embryos. This pathway may be important for maintaining genome stability in pericentric heterochromatin, proper nuclear organization, and other unexplored functions.

Our findings are consistent with a study that detected bi-directional transcription of the *1.688* satDNA family in ovaries (Usakin, et al. 2007) and a recent analysis of satDNA-derived piRNAs in RDC mutants (Chen, Luo, et al. 2020). Usakin et al. found that *1.688* transcript abundance is elevated in mutants of two piRNA processing genes, *spn-E* and *aub* (Usakin, et al. 2007), suggesting that *1.688* is targeted by piRNAs, similar to TEs. However, the origins of the *1.688* piRNAs and how the transcription of precursors is regulated was unclear (Usakin, et al. 2007). Here we provide evidence that most satellite-derived transcripts and small RNAs reported in previous studies (Aravin, et al. 2003; Saito, et al. 2006; Usakin, et al. 2007; Rosic, et al. 2014; Chen, Kotov, et al. 2020; Chen, Luo, et al. 2020) come from the heterochromatin-dependent transcription of the large satDNA blocks. The role of these piRNAs in ovaries remains unknown, and we understand even less about piRNA biogenesis and function in *D. melanogaster* testes, where we also detect satDNA-derived piRNAs. Proportionally, far fewer piRNAs in the male germline are derived from TEs than in the female germline (Nishida, et al. 2007; Nagao, et al. 2010; Quenerch’du, et al. 2016), suggesting roles outside of TE repression. For example, recent studies implicate piRNA pathways in intragenomic conflicts (*e.g*. male meiotic drive; Gell and Reenan 2013; Courret, et al. 2019), with satDNAs often at the center of these conflicts.

While it will take more work to understand the role of satDNA-derived transcripts in the germline, we hypothesize that the maternal deposition of these piRNAs contributes to heterochromatin establishment at satDNAs in the early embryo (Figure 5). Maternal deposition of Piwi contributes to heterochromatin establishment in the embryo (Gu and Elgin 2013), and Piwi-dependent H3K9me3 deposition at canonical piRNA clusters is important for subsequent piRNA production at piRNA clusters (Akkouche, et al. 2017). Similar to piRNA clusters, we found evidence that both H3K9me3 chromatin and piRNA production from complex satDNA is reduced when transiently depleting Piwi in the embryos, suggesting a role for the piRNA pathway in heterochromatin establishment at satDNA loci (Figure 5). We propose a simple model of self-regulation, where Piwi, guided by satDNA-derived piRNAs, establishes H3K9me3 at satDNA, marking the satDNAs as piRNA production sites later in development (Figure 5). While a contributor, Piwi might not be the only factor necessary for heterochromatin establishment in embryos (Wei, et al. 2021). And once established, the maintenance of heterochromatin at piRNA clusters and satDNAs is not Piwi-dependent (Table S10) (Klenov, et al. 2014; Mohn, et al. 2014). Therefore, the piRNA pathway is likely to be one of several factors important for proper packaging and regulation of repeat-rich regions of the genome (Pal-Bhadra, et al. 2004; Gu and Elgin 2013).

The consequences of disrupting satDNA packaging/regulation are likely to be complicated. The ramifications could be especially serious if a reduction in heterochromatin at satDNA in early embryos affects heterochromatin in all tissues (reviewed in Janssen, et al. 2018) and/or if establishing heterochromatin at satDNA loci serves as nucleation points for the larger-scale heterochromatinization of pericentric regions. First, heterochromatic regions form a distinct phase-separated nuclear compartment that contributes to nuclear organization and gene regulation (Larson, et al. 2017; Strom, et al. 2017), and chromocenter formation (Jagannathan, et al. 2018). Un-regulated satDNA may disrupt this organization (Novo, et al. 2020) and lead to cell death (Jagannathan, et al. 2019). Second, de-repressed satDNA may lead to genome instability (Peng and Karpen 2007) including chromosomal structural rearrangements (reviewed in Janssen, et al. 2018). In the short term, rearrangements involving satDNA may lead to mitotic defects in the developing embryo, as they can affect chromosome segregation (Ferree and Barbash 2009; Ferree 2014). Over longer evolutionary time scales, these rearrangements contribute to variation in satDNA organization between individuals and species, and may cause genetic incompatibilities between closely-related species (Ferree and Barbash 2009). SatDNAs are indeed among the most rapidly evolving sequences in genomes (reviewed in Ferree and Prasad 2012; Plohl, et al. 2012).

Many mysteries remain surrounding the functions of the piRNA pathway outside of its role in controlling TE activity. Our finding that the piRNA pathway regulates satDNA suggests a general role for the piRNA pathway and for maternal satDNA-derived RNAs in remodeling chromatin in the developing embryo. This initial establishment of heterochromatin may be an important step in ensuring genome integrity throughout development and in adult tissues, but this remains an open question. Moving forward, it will be important for piRNA studies to continue to focus on satDNA and how these dynamic compartments of the genome contribute to genome function and stability.

## MATERIALS AND METHODS

### *Drosophila* stocks

Iso-1 was used as the wild type strain, unless stated otherwise. In the qPCR validation experiment, *rhi* mutants (*rhi^-^*) are transheterozygotes from the Vienna Drosophila Resource Center (VDRC 313487 and 313488) as are the *moonshiner* mutants (*moon^-^*) (VDRC 313735 and 313738) as described in (Andersen, et al. 2017). Based on the origin and genetic background of these mutants, *w^1118^* or the progeny from OregonR (Ore) crossed to *w^1^* were used as the wild type controls for *rhi^-^* and *moon^-^*. All flies were maintained at 23°C on cornmeal medium.

### Small RNA-seq

Six- to eight-day-old testes were dissected in RNase free PBS buffer. Total RNA was extracted using *mir*Vana™ miRNA Isolation Kit (Ambion) with procedures for isolating RNA fractions enriched for small RNAs <200nt), then treated with RNase free DNase I (Promega) at 37°C for 1 hour. Library preparation and sequencing were performed by Genomics Research Center at University of Rochester. Briefly, 2S rRNA was depleted (Wickersheim and Blumenstiel 2013), small RNA library was prepared with TruSeq Small RNA Library Prep Kit (Illumina) and sequenced by Illumina platform HiSeq2500 Single-end 50 bp.

### Total RNA-seq

Six- to eight-day-old testes were dissected in RNase free PBS buffer. Total RNA was extracted using *mir*Vana™ miRNA Isolation Kit (Ambion) with procedures for isolating RNA fractions enriched for long RNAs (>200nt), then treated with RNase free DNase I (Promega) at 37°C for 1 hour. Library preparation and sequencing were performed by Genomics Research Center at University of Rochester. Briefly, rRNA was removed and total RNA library was prepared with TruSeq Stranded Total RNA Library Prep Human/Mouse/Rat (Illumina) and sequenced by Illumina platform HiSeq2500 Paired-end 125 bp.

### SatDNA analysis

Reads were mapped to the heterochromatin-enriched genome assembly (Chang and Larracuente 2019) and counted based on their annotations (*e.g. Rsp* or *1.688*). Due to the highly repetitive nature of satDNAs, around 80% of total RNA-seq and 99% of small RNA-seq reads that are mapped to satDNA regions are not uniquely assigned; discarding these multiple mapped reads would result in loss of statistical power in the satDNA analysis. To deal with this, multiple mapped reads were randomly assigned to one of their multiple best mapping locations, unless stated otherwise. Reads were then counted based on the annotations of their assigned mapping locations. Because there is high sequence similarity among the *1.688* subfamily repeats (*260-bp, 359-bp, 353-bp, 356-bp*), all *1.688* subfamilies were combined, unless stated otherwise. A similar approach was used in our analysis of piRNA clusters, except that only uniquely mapped reads were counted so that the published results could serve as controls for our method. Additional details specific to small RNA-seq, RNA-seq, ChIP-seq, and RIP-seq analyses are given below.

### RNA-seq analysis

All total RNA-seq datasets reanalyzed in our study are listed in Table S1. Total RNA-seq reads were trimmed for adaptors and then mapped to the genome using Bowtie2 (Langmead and Salzberg 2012). A customized python script was used to count reads that mapped to each repeat feature or piRNA cluster, and RPM values were reported by normalizing raw counts to 1,000,000 total mapped reads (https://github.com/LarracuenteLab/Dmelanogaster_satDNA_regulation (Wei 2020), htseq_bam_count_proportional.py). For the *1.688* subfamilies, all subfamilies were combined into one *1.688* category, although analyzing each by subfamily (e.g. *353-bp*, *356-bp*, *359-bp*, *260-bp*) does not change our conclusions (https://github.com/LarracuenteLab/Dmelanogaster_satDNA_regulation) (Wei 2020). For results shown in Table S8, DESeq2 (Love, et al. 2014) was used to perform differential expression analysis of the raw counts with combined data from different studies (Mohn, et al. 2014; Andersen, et al. 2017), with experimental condition and associated study as covariates. This analysis method is conservative and leads to smaller log2 fold changes than published results of piRNA clusters. For comparison with the published results, a similar approach was used to analyze piRNA clusters (Mohn, et al. 2014; Andersen, et al. 2017). Briefly, quantification of reads mapping to 1-kb windows inside each piRNA cluster was estimated using a customized python script (https://github.com/LarracuenteLab/Dmelanogaster_satDNA_regulation (Wei 2020), htseq_bam_count_proportional.py), and subsequent differential expression analysis between mutants and wildtype was done using DESeq2 (Love, et al. 2014; results shown in Figure 3–figure supplement 2).

### Small RNA-seq analysis

All small RNA-seq datasets reanalyzed in our study are listed in Table S1. Small RNA-seq reads were trimmed for adaptors, then mapped to the genome using Bowtie (Langmead 2010). A customized python script (https://github.com/LarracuenteLab/Dmelanogaster_satDNA_regulation; Wei 2020, htseq_bam_count_proportional.py) was used to count reads that mapped to each repeat feature or piRNA cluster. To control for differences in small RNA abundance and compare across samples, raw counts were then normalized to the number of reads that mapped to either miRNAs or the *flamenco* piRNA cluster. The difference in expression was represented by the log2 fold changes of these normalized counts in mutants compared to wild type (*i.e*., log2(count_mutant_/count_WT_)) for each repeat and piRNA cluster. We further analyzed the size distribution and relative nucleotide bias at positions along each satDNA by extracting reads mapped to the satDNA of interest using a customized python script (https://github.com/LarracuenteLab/Dmelanogaster_satDNA_regulation; Wei 2020, extract_sequence_by_feature_gff.py). The 10nt overlap Z-score of piRNAs mapped to each satDNA was calculated using piPipes (Han, et al. 2015). To determine which parts of these repeats are represented in piRNA or ChIP datasets, the read pileup patterns along the consensus sequence of a satDNA were examined (*e.g*. Figure 2–figure supplement 2). Reads (ChIP or piRNA) mapping to a particular satDNA or genomic satDNA variant (as a control) were BLAST-ed to the consensus dimer (for *1.688* satellite) or trimer (for *Rsp* because it has left and right consensus sequences), and then coordinates were converted along a dimer/trimer to coordinates along a monomer/dimer consensus sequence. All plots were made in R (R core team 2017).

### ChIP/RIP-seq analysis

All total ChIP-seq and RIP-seq datasets reanalyzed in our study are listed in Table S1. ChIP-seq and RIP-seq reads were trimmed for adaptors and mapped to the genome using Bowtie2 (Langmead and Salzberg 2012). A customized python script (https://github.com/LarracuenteLab/Dmelanogaster_satDNA_regulation; Wei 2020, htseq_bam_count_proportional.py) was used to count reads that mapped to each repeat feature or piRNA cluster. Raw counts were normalized to 1,000,000 total mapped reads.

For the ChIP-seq results, enrichment scores of each repeat and piRNA cluster were reported by comparing the ChIP sample with the antibody of interest to its no-antibody input control sample. For ChIP-seq analyses, consider satDNA as discrete loci rather than repeat unit types is appropriate because some loci are composed of several repeat types. To examine the large blocks of heterochromatic satDNA chromatin for the Rhi and H3K9me3 ChIP-seq analyses, euchromatic *1.688* satDNAs were excluded and only reads that map uniquely to satDNA loci were analyzed. Heterochromatic satDNA loci were defined as discrete loci on chromosomes: 2L (2L_2: 402701-460225; the *260* locus), 3L (3L_3: 46695-106272; primarily *353-bp* and *356-bp* repeats), and the unmapped contigs (Contig101 and Contig9; *353-bp, 356-bp*, and *359-bp* repeats). Our conclusions do not change when we look at all reads (not just uniquely mapped; https://github.com/LarracuenteLab/Dmelanogaster_satDNA_regulation; Wei 2020). These analyses were repeated by combining all *1.688* subfamilies into a single category, and each subfamily was analyzed separately (*e.g*. all *353-bp* repeats combined) but the conclusions do not change (https://github.com/LarracuenteLab/Dmelanogaster_satDNA_regulation; Wei 2020). Euchromatic controls are included for the Rhi and H3K9me3 ChIP-seq analyses. Here, the euchromatic control correspond to the median enrichment score for protein coding genes that are 5Mb distal from heterochromatin boundaries (Riddle, et al. 2011) and piRNA clusters.

For the RIP-seq analyses, reported was the percentage of reads mapped to each repeat and piRNA cluster with miRNAs as the negative control. For the *1.688* subfamilies, all subfamilies were combined into one *1.688* category, although analyzing each by subfamily (e.g. *353-bp, 356-bp, 359-bp, 260-bp*) does not change the conclusions (https://github.com/LarracuenteLab/Dmelanogaster_satDNA_regulation; Wei 2020).

### RNA FISH

A Cy5-labeled oligo probe (5’-Cy5TTTTCCAAATTTCGGTCATCAAATAATCAT-3’) previously described in (Ferree and Barbash 2009) was used to detect *1.688* transcripts from all subfamilies except *260-bp* on chromosome *2L*. Custom Stellaris FISH Probes were designed for *Rsp* (Table S11), and RNA FISH was performed following the manufacturer’s instructions (Biosearch Technologies, Inc). Three- to six-day-old ovaries and testes were dissected in RNase free PBS buffer, fixed with 4% paraformaldehyde in PBS buffer at room temperature for 30 minutes, and then washed twice with PBS for 5 minutes. To permeabilize, tissues were kept in RNase free 70% ethanol at 4°C overnight. The ethanol was aspirated, and samples washed with Stellaris wash buffer on a nutating mixer for 3 minutes and kept still for 2 minutes at room temperature. Hybridization was then performed with each probe in Stellaris hybridization buffer in a humidity chamber at 37°C overnight. The working concentration was 100nM for the oligo probe and 125nM for the Stellaris probes. From this point, samples were kept in the dark. The samples were washed with Stellaris wash buffer twice at 37°C for 30 minutes each. Samples were then transferred to mounting medium containing DAPI and imaged with Leica SP5 laser scanning confocal microscope.

For RNaseA controls, after fixation and permeation, tissues were treated with RNase A (100ug/ml) in RNase digestion buffer (5 mM EDTA, 300 mM NaCl, 10 mM Tris-HCl pH 7.5, cold spring harbor protocols, http://cshprotocols.cshlp.org/content/2013/3/pdb.rec074146.full) at 37°C for 1hr and washed three times with Stellaris wash buffer at room temperature for 10 minutes before hybridization.

For RNase H controls, after probe hybridization and washing, tissues were treated with 1.5uL RNase H (5,000 units/ml; New England Biolabs) in 50uL final volume in 1X RNAse H buffer at 37°C for 2 hours and washed three times with Stellaris wash buffer at room temperature for 10 minutes before mounting and imaging.

### qPCR

For genomic DNA qPCR, three- to eight-day-old flies were mashed with pipette tips for 5-10 seconds and incubated in buffer (10 mM Tris-Cl pH 8.2, 1 mM EDTA, 25 mM NaCl, 200 ug/ml Proteinase K) at 37°C for 30 minutes (Gloor and Engels 1992). To extract nucleic acids, a mixture of phenol/Sevag (1:1) of equal volume was added, and the samples vortexed for 45 to 60 seconds and then centrifuged for 3-5 minutes. The aqueous top layers were saved, an equal volume of Sevag added, and the samples vortexed for 30 seconds then centrifuged for 1 minute. The aqueous top layers were saved and a second Sevag extraction performed. Diluted nucleic acid samples (concentration of 0.04ng/uL) were used for qPCR to determine the repeat copy numbers in the genome. Repeat copy numbers are normalized to the tRNA:Lys-CTT copy numbers.

For RNA qRT-PCR, three- to six-day-old ovaries were dissected in RNase free PBS buffer, and total RNA was extracted using the *mir*Vana™ miRNA Isolation Kit (Ambion). RNA samples were treated with RNase free DNase I (Promega) at 37°C for 1 hour. The RNA samples were reverse transcribed using random hexamer primers and M-MuLV Reverse Transcriptase (New England Biolabs) and the resulting cDNA subjected to qPCR. To exclude the possibility of DNA signal in qRT-PCR experiments, controls with no Reverse Transcriptase enzyme were used for all samples in the reverse transcription step. Expression levels were normalized to ribosomal protein S3 (RPS3) expression. To detect the transcript abundance difference between wild type and mutant, ΔΔCT was calculated (Livak and Schmittgen 2001).

The replicate number for genomic DNA qPCR is 2-4, and for RNA qRT-PCR is 4-6. The sequences of primers used are: *Rsp* (forward: GGAAAATCACCCATTTTGATCGC, reverse: CCGAATTCAAGTACCAGAC); tRNA (forward: CTAGCTCAGTCGGTAGAGCATGA, reverse: CCAACGTGGGGCTCGAAC); RPS3: (forward: AGTTGTACGCCGAGAAGGTG, reverse: TGTAGCGGAGCACACCATAG).

### Northern Blot Analysis

Isolation of total RNA and RNase controls: stocks of *D. melanogaster* were chosen which represented a range of *Rsp* repeat copy numbers; flies were collected (0 to 20 hours old) and aged for six days. Ovaries were dissected from approximately 20 females (i.e., 6.0 to 6.8 days old) from each stock, and total nucleic acid isolated using a standard phenol/Sevag procedure (Khost, et al. 2017). Total nucleic acid was then treated with DNase I as recommended (20 units; Promega), re-extracted with phenol/Sevag, and ethanol precipitated. Total RNA was resuspended in distilled water. The integrity of the RNA was checked on 1% agarose gels, and the concentration estimated by an optical density at 260 nm.

For RNase controls, 10 μg of total RNA was resuspended in 50 mM NaCl, 5 mM EDTA, 10 mM Tris pH 7.5, 100 μg/ml RNaseA and incubated at 37°C for 30 minutes. Samples were phenol/Sevag extracted, 10 μg of ytRNA added as carrier, and ethanol precipitated.

Northern Blot Analysis: Total RNA (10 μg)/ RNase controls were suspended in 1×MOPS [0.04 M MOPS (morpholinepropanesulfonic acid) pH 7.0, 0.01 M Na acetate, 0.001 M EDTA], 2.2 M formaldehyde, 50% formamide. The RNA was then heated at 65°C for 15 min, placed on ice, and one-tenth volume loading buffer (1× MOPS, 50% formamide, 2.2 M formaldehyde, 4% Ficoll, 0.25% bromophenol blue) added. RNAs were separated on a 1% agarose gel containing 0.5 M formaldehyde/1 × MOPS at 40 V for 3 h. Standard RNA lanes were cut from the gel and stained with ethidium bromide to monitor electrophoresis. Gels were washed for 25 minutes in sterile water (with 4 changes). RNA was transferred to GeneScreen Plus nylon membrane (prewet in 10× SSC) by capillary action using 10× SSC. After transfer, the nylon membrane was rinsed in 2× SSC, UV crosslinked, and then baked for 2 h under vacuum at 80°C. The membrane was prehybridized in 2× SSC, 5× Denhardt’s solution, 1% sodium dodecyl sulfate (SDS), 10% polyethylene glycol (PEG-molecular weight, 8,000), 25 mM sodium phosphate (pH 7.2), 0.1% sodium pyrophosphate, and 50% formamide for 3 h at 55°C. Hybridizations were done overnight at 55°C in the same buffer containing a biotinylated RNA probe (see slot blot; primers: T7_rsp2 5’-TAATACGACTCACTATAGGGCCGAATTCAAGTACCAGAC-3’ and rsp1 5’-GGAAAATCACCCATTTTGATCGC-3’). The hybridized membranes were washed in 1M sodium phosphate pH 6.8, 0.5M EDTA, 5% SDS (2X, 10 minutes each) at 60°C and then at 1M sodium phosphate pH 6.8, 0.5M EDTA, 1% SDS (3X, 10 minutes each) at 65°C. The washed membranes were then processed as recommended for the Chemiluminescent Nucleic Acid Detection Module (ThermoScientific), and the signal recorded on a ChemiDoc XR+ (Bio-Rad).

### Slot blot

Five female flies were mashed and the total nucleic acid phenol/Sevag extracted as described above for qPCR. Approximately 200 ng of the nucleic acid was denatured (final concentration 0.25 M NaOH, 0.5 M NaCl) for 10 minutes at room temperature, the sample transferred to a tube with an equal volume of ice-cold loading buffer (0.1X SSC, 0.125 M NaOH) and left on ice. The slot blotter was then prepared and samples loaded as recommended for the 48-well BioDot SF microfiltration apparatus (Bio-Rad). After loading, the wells were washed with 200 μl of loading buffer. The nylon membrane (GeneScreen Plus) was then rinsed for 2 minutes with 2X SSC before being UV crosslinked (Stratalinker). The membrane was first hybridized with a biotinylated rp49 RNA probe in North2South hybridization solution (ThermoScientific) at 65°C overnight. The membrane was processed as recommended for the Chemiluminescent Nucleic Acid Detection Module (ThermoScientific), and the signal recorded on a ChemiDoc XR+ (Bio-Rad). The membrane was then stripped with a 100°C solution of 0.1X SSC/ 0.5% SDS (3 times for ~20 minutes each) and re-hybridized with a *Rsp* probe (60°C overnight) and processed as above. Signals were quantitated using the ImageLab software (Bio-Rad). We determined the relative signal compared to Iso-1 for each line (5-7 replicates), and then estimate the *Rsp* copy number by scaling the relative slot blot signal to our estimate of *Rsp* copy number in Iso-1 (1100 repeats). Our Iso-1 estimate is based on *Rsp* count in a long-read assembly, which is supported by empirical slot blots (Khost, et al. 2017).

To make the biotinylated RNA probes, gel extracted PCR amplicons (primers: *Rsp* 5’-TAATACGACTCACTATAGGGGAAAATCACCCATTTTGATCGC-3’ and 5’-CCGAATTCAAGTACCAGAC-3’; rp49 5’-TAATACGACTCACTATAGGGCAGTAAACGCGGTTCTGCATG-3’ and 5’-CAGCATACAGGCCCAAGATC-3’) were transcribed using the Biotin RNA Labeling Mix (Roche) and T7 polymerase (Promega).

## Acknowledgements

This work was supported by the National Institutes of Health General Medical Sciences (R35 GM119515 to AML), a Stephen Biggar and Elisabeth Asaro fellowship in Data Science to AML, and a University of Rochester Messersmith Fellowship to XW. We thank Drs. Ching-Ho Chang, John Sproul, Cécile Courret, and Lucas Hemmer for providing feedback on the manuscript. We also thank the University of Rochester Center for Integrated Research Computing for access to computing facilities and the University of Rochester Genomics Research Center for sequencing services.

## Data availability

Sequencing data generated in this paper are available in the NCBI Sequence Read Archive under project accession PRJNA647441. All data files and code to recreate analyses and figures are deposited in GitHub (https://github.com/LarracuenteLab/Dmelanogaster_satDNA_regulation; Wei 2020) and at the Dryad Digital Repository (link forthcoming).

