## Supplemental figures for "Heterochromatin-dependent transcription of satellite DNAs in the *Drosophila melanogaster* female germline"

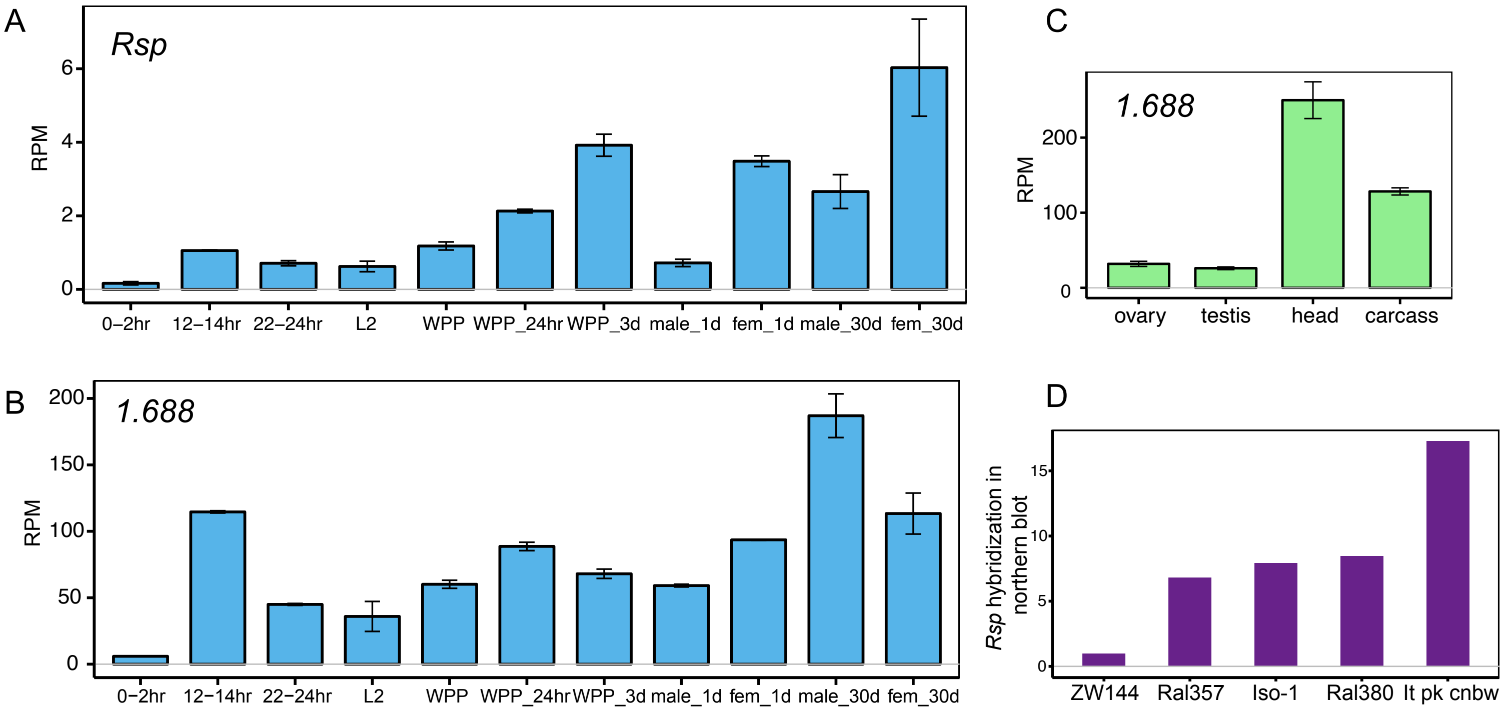


Figure 1–figure supplement 1. SatDNAs are expressed across tissues and developmental stages. (A-B) SatDNA transcription level in various developmental stages (A for *Rsp* and B for *1.688*) . L2: 2nd instar larvae. WPP: white prepupae. Fem: female. Complex satDNA expression increases with adult age. (C) *1.688* satDNA transcription level in various tissues (corresponding result for *Rsp* is shown in Figure 1A). Carcass: whole body without the head, reproductive organs and digestive tract. (D) Northern signal quantification of *Rsp* transcript abundance. Data for A, B and C from modENCODE (Graveley, et al. 2011; Brown, et al. 2014).

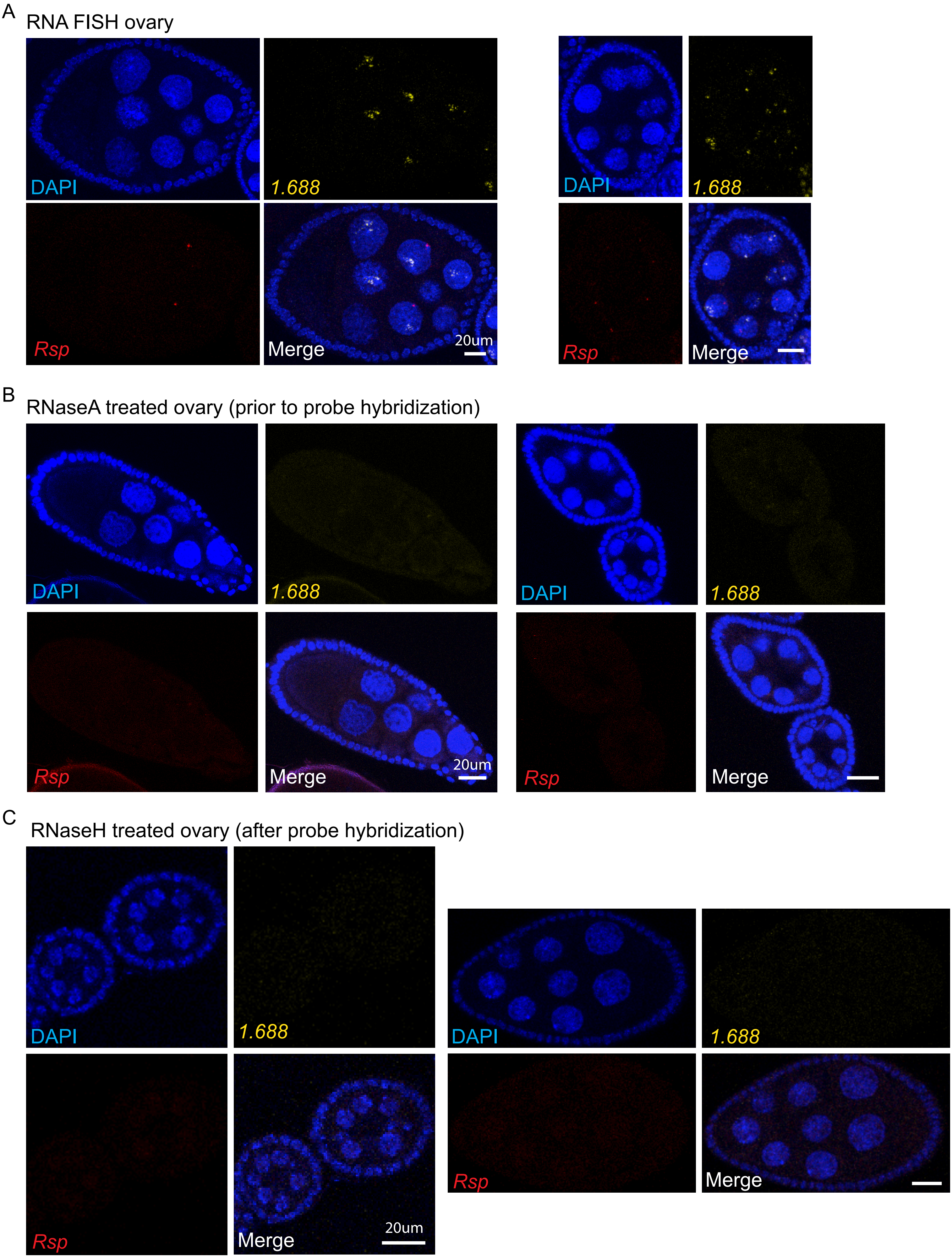


Figure 1–figure supplement 2. RNA FISH signals of satDNAs are from RNAs, not DNAs. RNA FISH of *Rsp* and *1.688* in ovary (A), with RNaseA treatment prior to probe hybridization (B) and with RNaseH treatment after DNA probe hybridization (C). The *1.688* probe recognizes major *1.688* loci on chromosomes X and 3.

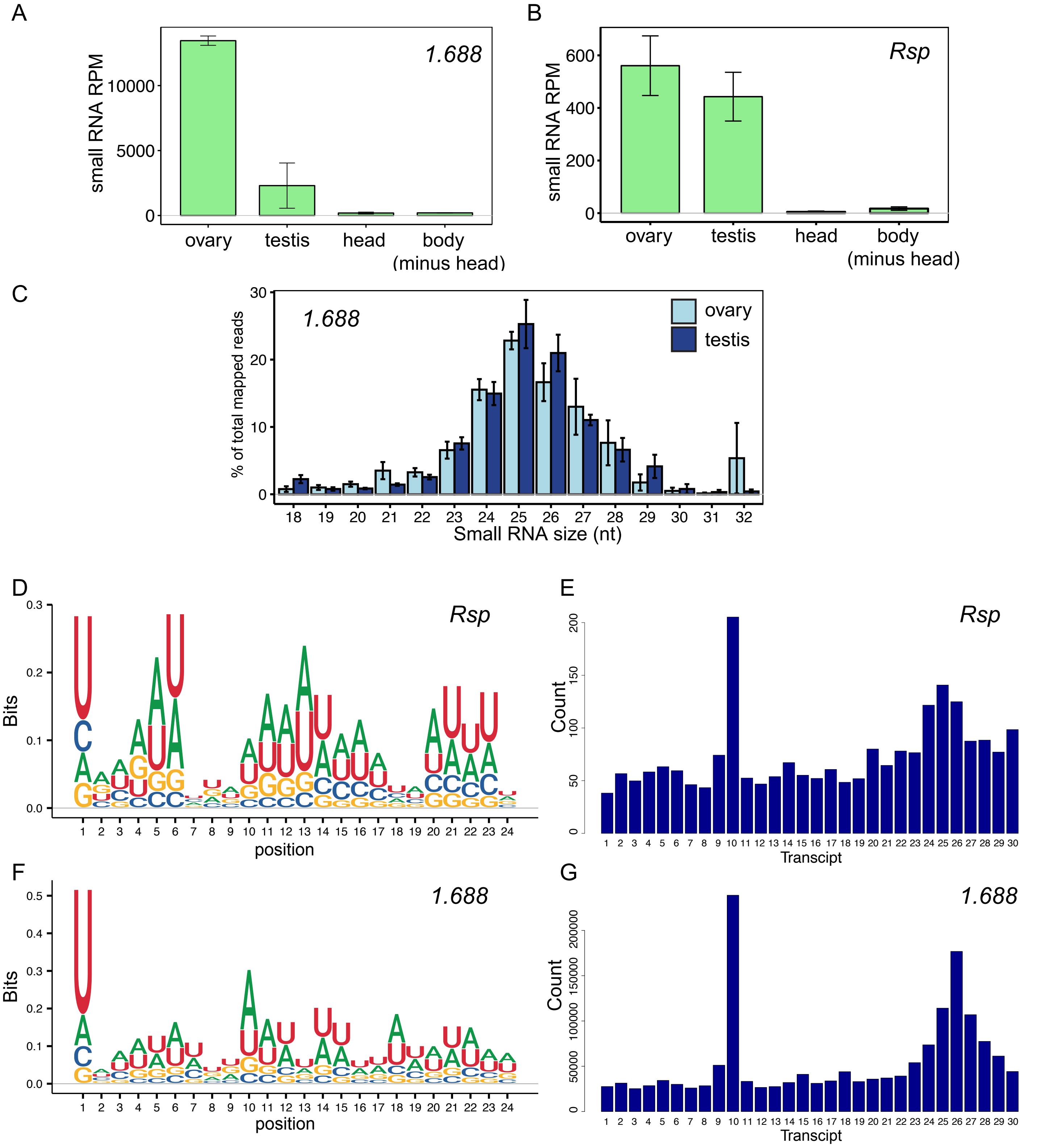


Figure 2–figure supplement 1. Satellite-derived RNAs are mainly processed into piRNAs in the germline. (A-B) Small RNA levels (RPM) in ovary, testis, head and body (minus head) for *1.688* (A) and *Rsp* (B). (C) Size distribution of *1.688* small RNAs in gonads. (D and F) Relative nucleotide bias of each position in small RNAs from *Rsp* (D) and *1.688* (F) satellite in ovary. (E and G) 5'-to-5' end distance analysis results of small RNAs from *Rsp* (E) and *1.688* (G) satellite in ovary show bias of 10-nt overlap. Z-score=4.55 for *Rsp* and 6.85 for *1.688* satellite. Data from (Ghildiyal, et al. 2010; Rozhkov, et al. 2010; Fagegaltier, et al. 2014; Mohn, et al. 2014; Quenerch'du, et al. 2016; Andersen, et al. 2017; Parhad, et al. 2017).

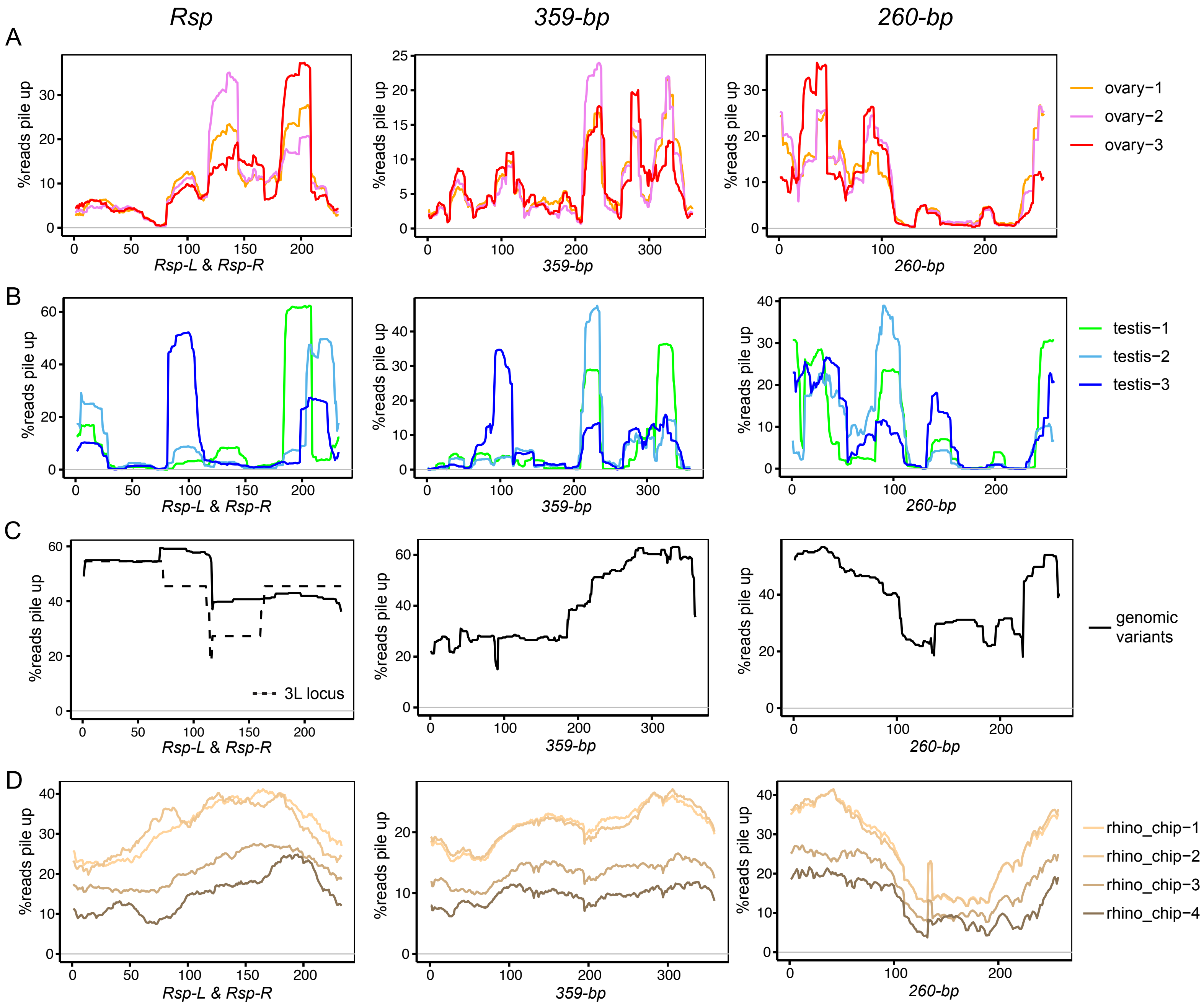


Figure 2–figure supplement 2. Non-uniform distribution of piRNA reads along satDNA consensus sequences. Small RNA reads pileup along *Rsp*, *260-bp* and *359-bp* (*1.688* satellite) consensus sequences for ovary (A) and testis (B). *Rsp-L* is 1-116nt and *Rsp-R* is 117-223nt in a *Rsp* dimer sequence. (C) Alignment depth of genomic satDNA repeat variants extracted from the genome assembly along consensus sequences. *Rsp* variants resident inside the *Ago3* intron on chromosome 3L (3L locus) is shown separately in dashed line. (D) Rhino ChIP-seq reads pileup along the satellite consensus sequences. Data from (Rozhkov, et al. 2010; Mohn, et al. 2014; Quenerch'du, et al. 2016; Andersen, et al. 2017; Parhad, et al. 2017; Zhang, et al. 2014).

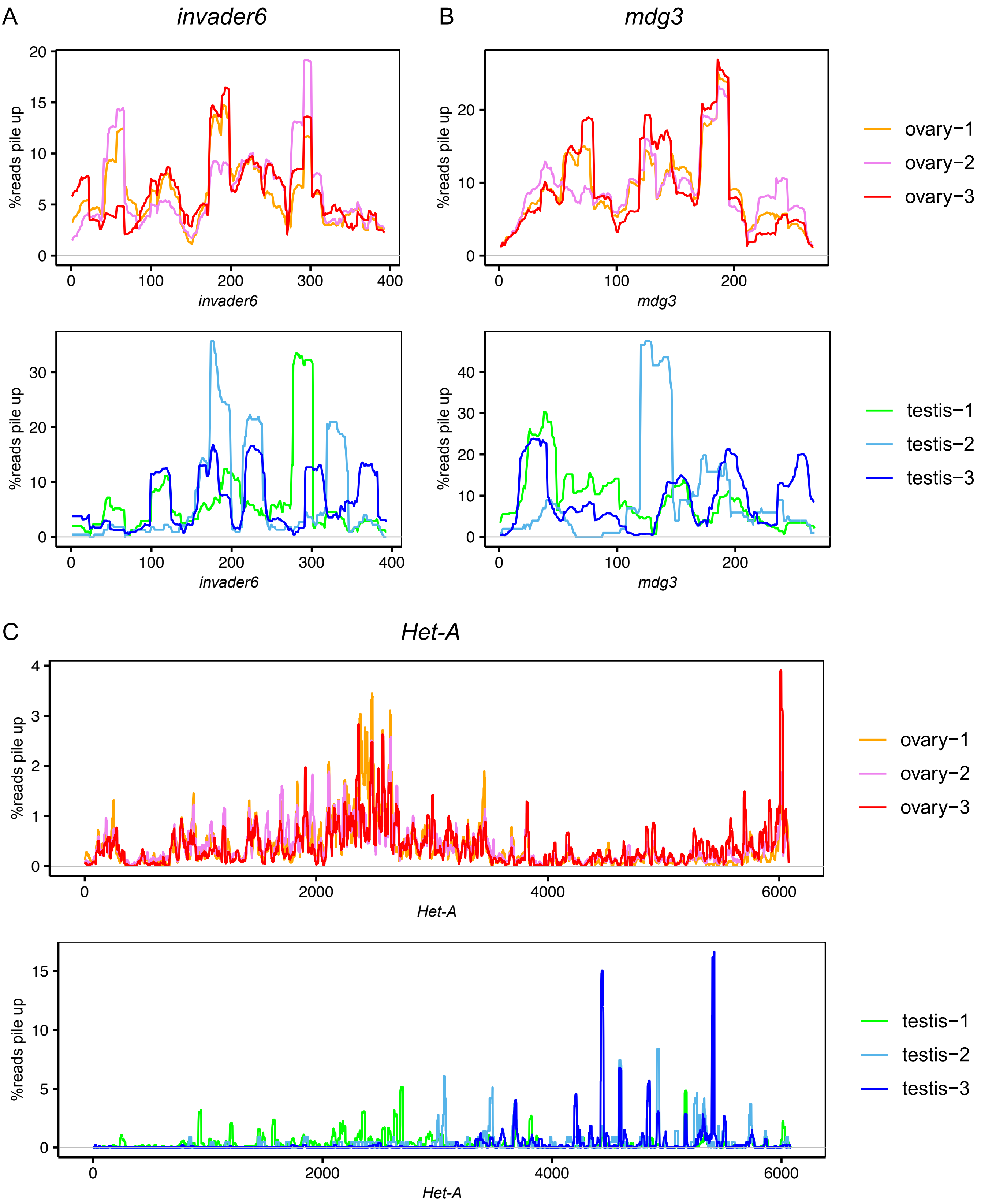


Figure 2–figure supplement 3. Non-uniform distribution of piRNA reads along the germline-dominant TE consensus sequences. Small RNA reads pileup along *Invader6* (A), *mdg3* (B) and *Het-A* (C) consensus sequences for ovary and testis. Data from (Rozhkov, et al. 2010; Mohn, et al. 2014; Quenerch'du, et al. 2016; Parhad, et al. 2017; Andersen, et al. 2017)

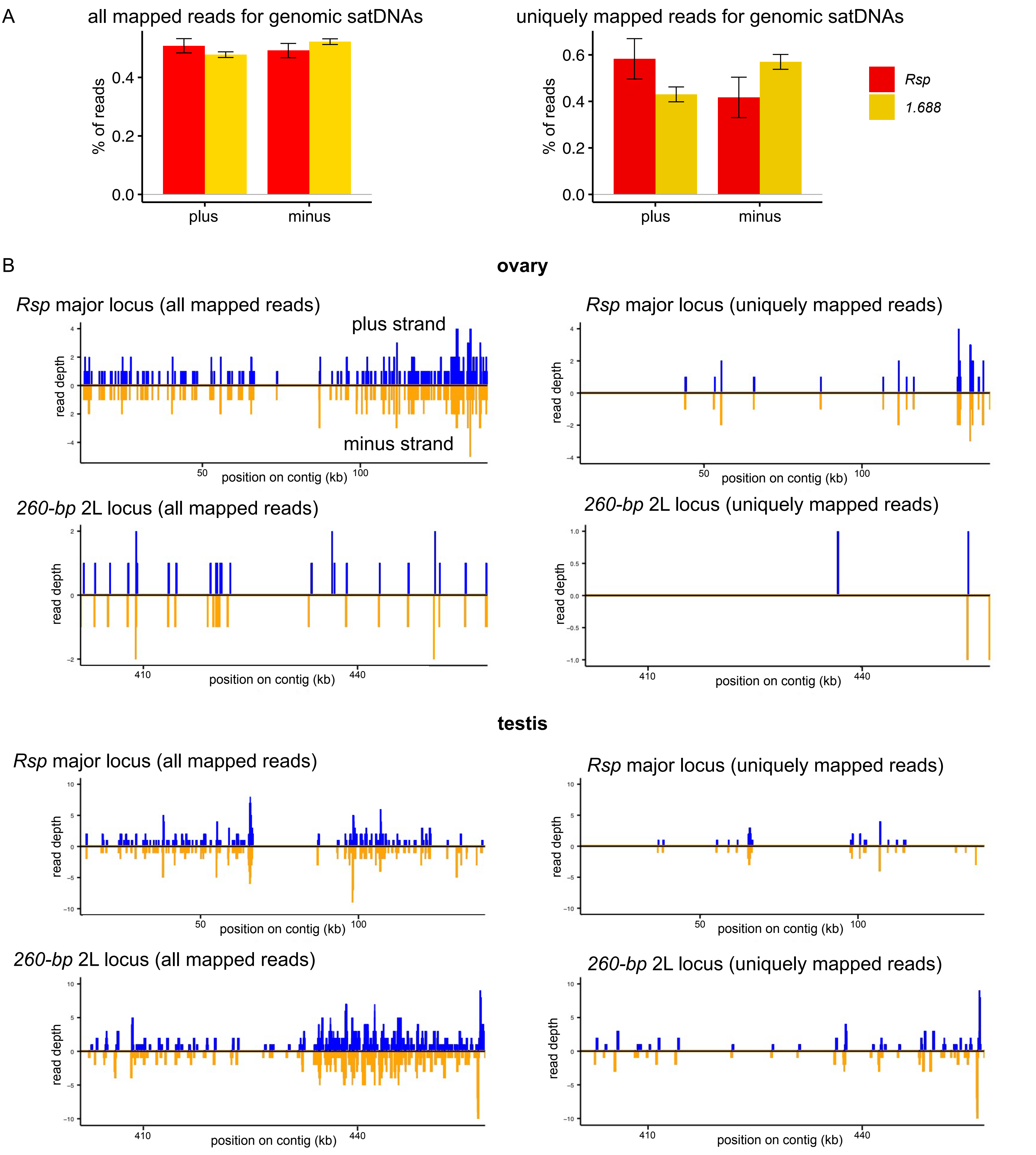


Figure 2–figure supplement 4. SatDNAs are transcribed from both strands. (A) Percentages of reads mapped to plus and minus strands at all genomic copies of *Rsp* or *1.688* satellites, for all mapped reads (left) and uniquely mapped reads (right). (B) Read depth for the plus and minus strands on the contigs containing the *Rsp* 2R major locus and *260-bp* 2L locus (Khost et al. 2017). Read depth of all reads mapped to each locus is shown on the left, and depth of uniquely mapped reads is shown on the right. Ovary data is from (Mohn, et al. 2014; Andersen, et al. 2017), and testis data is from this study.

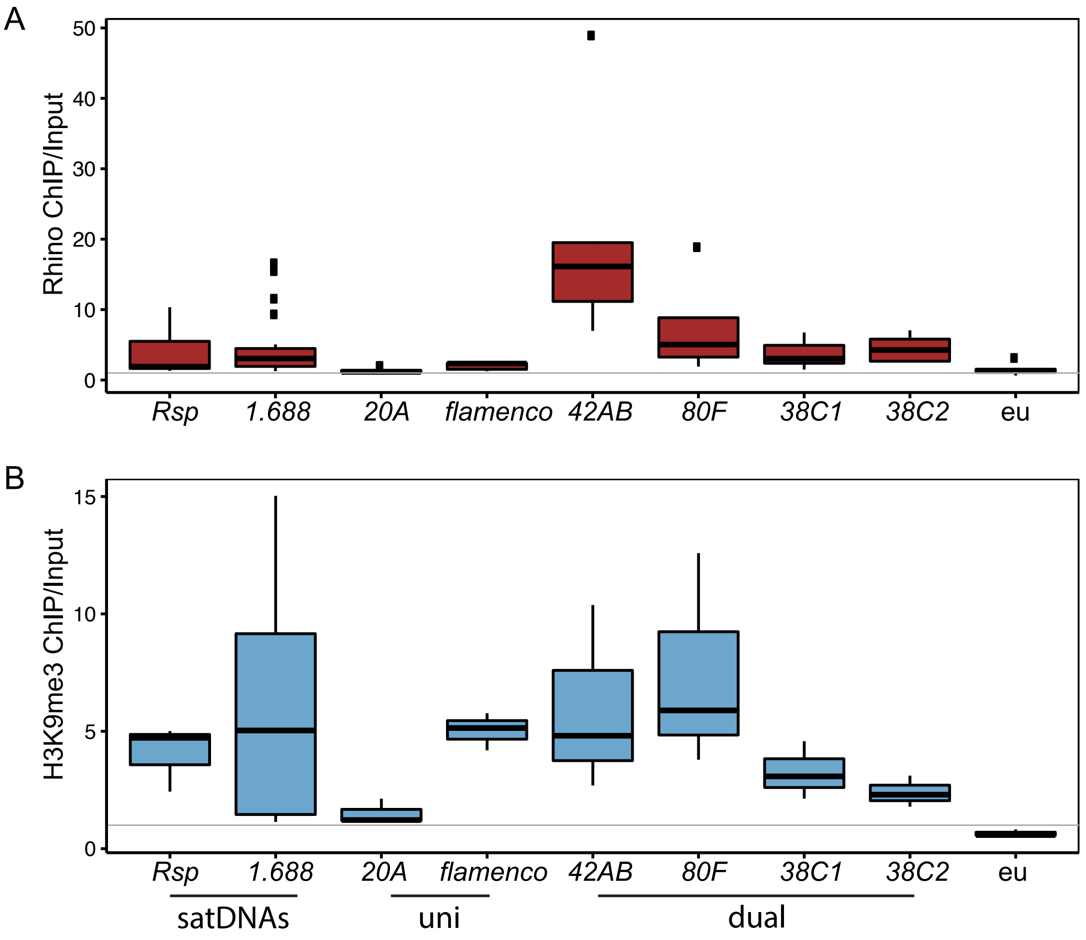


Figure 2–figure supplement 5. ChIP-seq result shows the chromatin state of satDNAs, uni-strand piRNA clusters, dual-strand piRNA clusters and euchromatin (eu). (A) Rhino ChIP-seq enrichment scores for *Rsp*, *1.688* (based on an analysis of discrete heterochromatic loci, details in MATERIALS AND METHODS), piRNA clusters, and euchromatin indicate that satDNAs are enriched for Rhino, resembling dual-strand piRNA clusters. Uni-strand piRNA clusters and euchromatin are not Rhino enriched. The Rhino enrichment results for *1.688* with all subfamilies across the genome are similar. (B) H3K9me3 ChIP-seq enrichment scores indicate that *Rsp*, *1.688* (based on an analysis of discrete heterochromatic loci, details in MATERIALS AND METHODS) and most piRNA clusters are enriched for H3K9me3, while euchromatin is not. Data from (Parhad, et al. 2017; Klenov, et al. 2014; Le Thomas, et al. 2014; Mohn, et al. 2014; Zhang, et al. 2014).

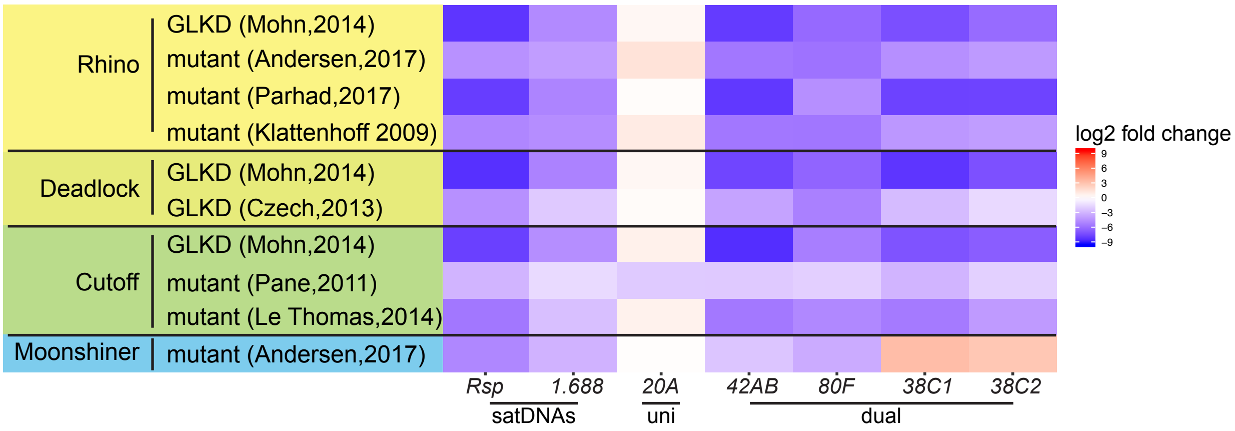


Figure 3-figure supplement 1. SatDNA loci are regulated by the heterochromatin-dependent transcription machinery in Drosophila ovaries. Heatmap showing the quantification of piRNA abundance from mutants of *rhino*, *cutoff*, *deadlock*, and *moonshiner* for satDNAs and piRNA clusters, normalized to the *flamenco* piRNA cluster. GLKD: germline knockdown. Complete list of log2 fold changes in Table S7. Data from (Klattenhoff, et al. 2009; Pane, et al. 2011; Czech, et al. 2013; Le Thomas, et al. 2014; Mohn, et al. 2014; Andersen, et al. 2017; Parhad, et al. 2017).

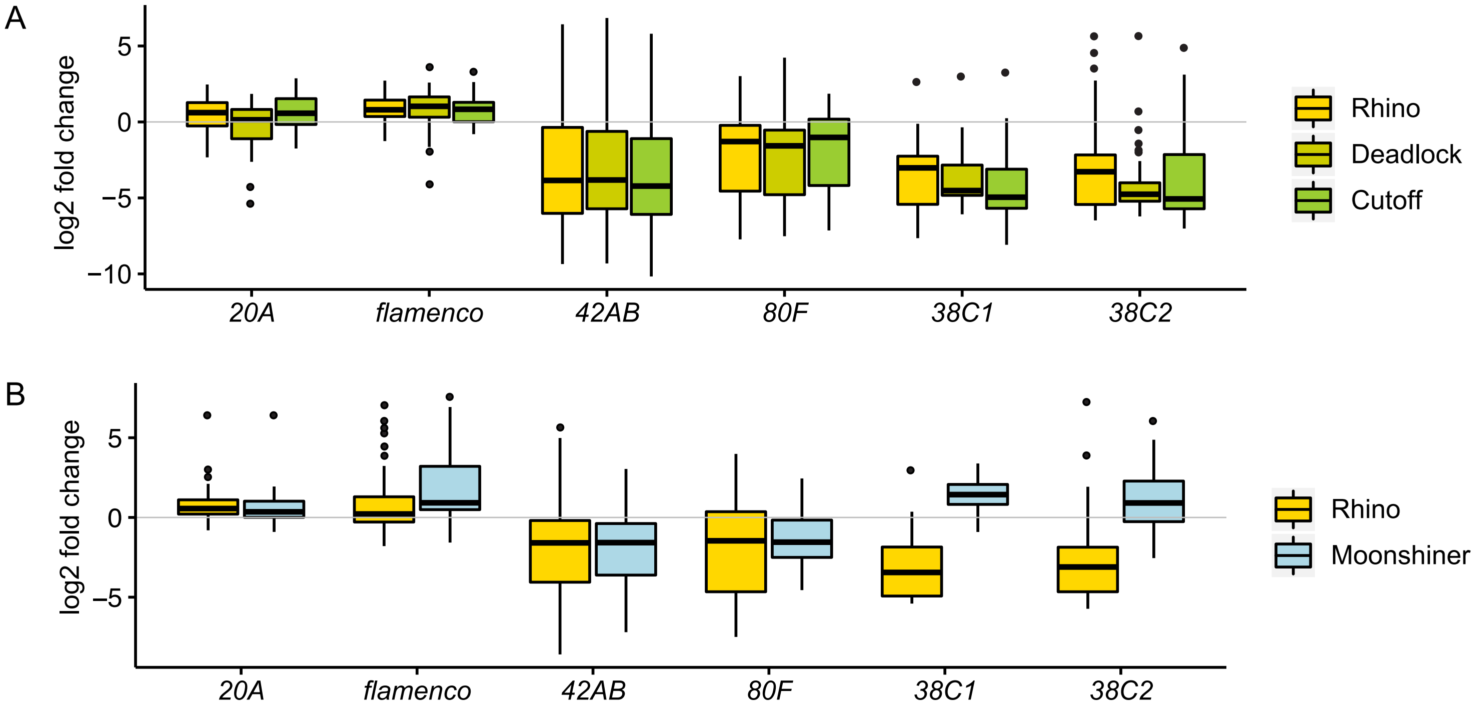


Figure 3-figure supplement 2. RDC and Moon mutants affect piRNA precursor transcription at piRNA clusters. Boxplots showing the quantification of total RNA abundance for piRNA clusters in RDC (A) and Rhi/Moon (B) mutant ovaries relative to wild type (log2 fold change of 1-kb windows). *20A* and *flamenco* are uni-strand piRNA clusters; *42AB*, *80F* and *38C1/2* are dual-strand piRNA clusters. Data from (Mohn, et al. 2014) for (A) and (Andersen, et al. 2017) for (B). A sliding window method is not feasible for analyzing satDNAs because it relies on uniquely mapped reads, of which there are relatively few at satDNA loci. Therefore, we instead count reads mapping to all genomic repeat variants for each satDNA and piRNA cluster (the whole locus; Table S8). The locus-wide results are comparable with the sliding window analysis, but more conservative.

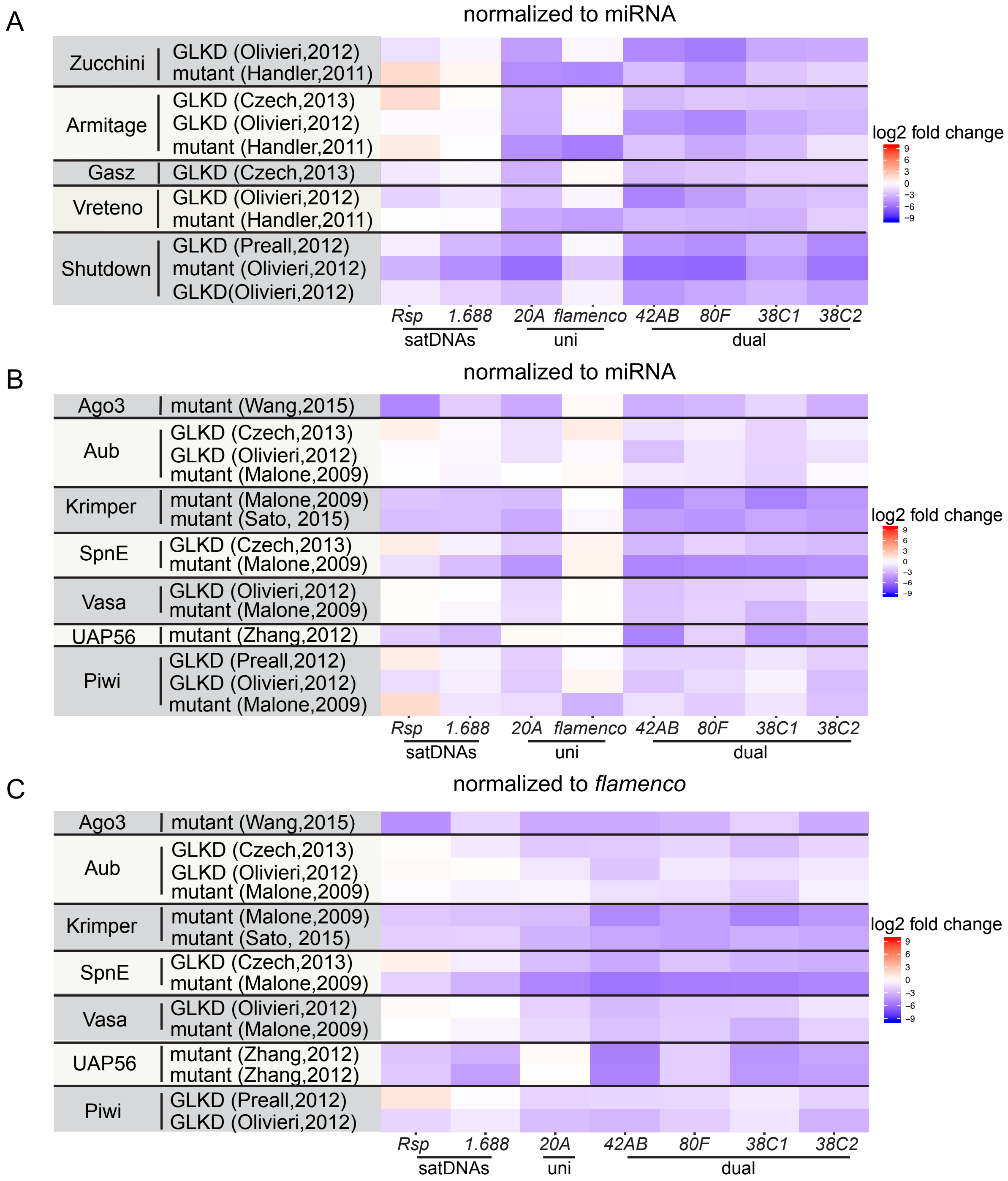


Figure 3-figure supplement 3. SatDNA piRNA production is affected in mutants of pathways involving piRNA precursor export, primary piRNA biogenesis, and the ping-pong cycle. Heatmap showing the quantification of small RNA abundance from mutants of proteins in the primary piRNA pathway (A), pathway for piRNA precursor export from the nucleus and ping-pong pathway (B-C). The results are consistent between replicates/studies about Vreteno, Shutdown, Krimper and UAP56, suggesting that complex satDNA piRNA production is regulated by these proteins. For some mutants including Aub, SpnE, and Piwi, results differed between complex satDNAs — *e.g.* the *1.688* family of satDNAs show consistently decreased piRNA levels, but not *Rsp* for Piwi. For other proteins such as Zucchini and Armitage, the patterns of change for satDNAs are variable between different datasets — *e.g.* satDNAs show decreased piRNA levels in one of the datasets, but increased levels in the other dataset for Zucchini. For all the datasets analyzed, all piRNA clusters behave as reported previously. GLKD: germline knockdown. *Flamenco* is only expressed in somatic tissues, GLKD doesn't affect its abundance. piRNAs are resistant to oxidation, for one of the datasets from (Zhang, et al. 2012), they were selected for sequencing after oxidation to exclude other types of small RNAs (*e.g.* siRNAs, miRNAs), so normalization to miRNA is not appropriate, thus it is excluded in (B). In *piwi* mutant from (Malone, et al. 2009), *flamenco* expression is affected and not appropriate as control for normalization, thus it is excluded in (C). Data from (Malone, et al. 2009; Handler, et al. 2011; Olivieri, et al. 2012; Preall, et al. 2012; Zhang, et al. 2012; Czech, et al. 2013; Sato, et al. 2015; Wang, et al. 2015).

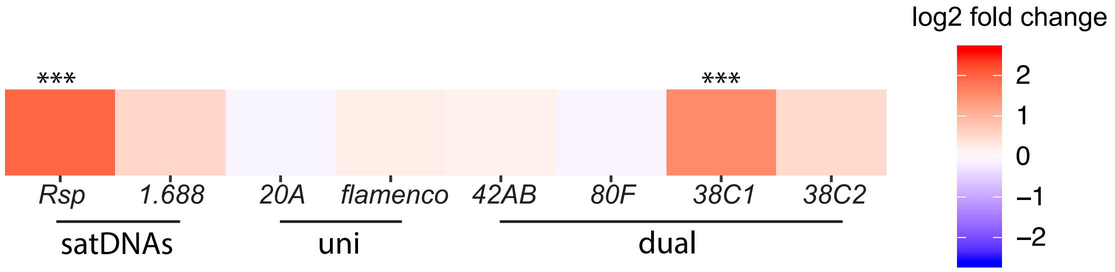


Figure 4–figure supplement 1. Log2 fold change of total RNA abundance shows satDNA long RNA levels increase in *piwi* embryonic knockdown ovaries compared to control. Adjusted p-values are reported by DESeq2. *20A* and *flamenco* are uni-strand piRNA clusters (uni), *42AB*, *80F* and *38C1/2* are dual-strand piRNA clusters (dual). *** adjusted p-value<0.001. Data from (Akkouche, et al. 2017).
