## Supplementary material for "Heterochromatin-dependent transcription of satellite DNAs in the *Drosophila melanogaster* female germline": TableS2

Table S2. Count in RPM values of *Rsp*/*1.688* transcripts in total and poly-A RNA-seq datasets from various tissues (data from modENCODE (Graveley, et al. 2011; Brown, et al. 2014))

| Repeat | RNA-seq type | Replicate | ovary | testis | carcass | head |
| --- | --- | --- | --- | --- | --- | --- |
| *Rsp* | Total RNA | 1 | 1.74 | 0.46 | 1.22 | 1.56 |
|  |  | 2 | 1.46 | 1.5 | 0.97 | 1.57 |
|  |  | 3 | 1.54 | 0.25 | 0.86 | 1.36 |
|  |  | 4 |  |  |  | 2.54 |
|  |  | 5 |  |  |  | 2.3 |
|  |  | 6 |  |  |  | 1.61 |
|  | polyA RNA | 1 | 0.17 | 0.01 | 0 | 0 |
|  |  | 2 | 0.13 | 0 | 0.01 | 0 |
|  |  | 3 |  | 0 |  | 0 |
|  |  | 4 |  |  |  | 0.03 |
|  |  | 5 |  |  |  | 0.08 |
| *1.688* | Total RNA | 1 | 37.17 | 25.09 | 137.87 | 191.56 |
|  |  | 2 | 33.08 | 29.56 | 122.89 | 211.17 |
|  |  | 3 | 25.42 | 23.55 | 124.27 | 228.41 |
|  |  | 4 |  |  |  | 217.12 |
|  |  | 5 |  |  |  | 316.69 |
|  |  | 6 |  |  |  | 333.75 |
|  | polyA RNA | 1 | 2.3 | 3.31 | 1.47 | 3.97 |
|  |  | 2 | 1.3 | 2.72 | 2.33 | 0.95 |
|  |  | 3 |  | 4.25 |  | 0.97 |
|  |  | 4 |  |  |  | 9.41 |
|  |  | 5 |  |  |  | 6.04 |
