## Supplementary material for "Heterochromatin-dependent transcription of satellite DNAs in the *Drosophila melanogaster* female germline": TableS3

Table S3. Slot blot estimate of Rsp copy number

| **fly strains** | **replicates** | | | | | | | **average** | **estimate copy number** |
| --- | --- | --- | --- | --- | --- | --- | --- | --- | --- |
|  | **1** | **2** | **3** | **4** | **5** | **6** | **7** |  |  |
| ZW144 | 0.2 | 0.21 | 0.27 | 0.22 | 0.15 | 0.25 |  | 0.2167 | 200 |
| Ral357 | 0.69 | 0.65 | 0.35 | 0.58 | 0.64 |  |  | 0.5820 | 600 |
| Ral380 | 2.74 | 2.29 | 1.9 | 1.95 | 1.75 |  |  | 2.1260 | 2300 |
| lt pk cn bw | 5.46 | 5.19 | 2.26 | 1.78 | 4.12 | 3.6 | 3.59 | 3.7143 | 4100 |

replicates: ratio of *Rsp* signal over *rp49* signal relative to that of Iso-1

estimate copy number: average ratio multiplied by the 1100 *Rsp* copy estimate for Iso-1 (Khost et al., 2017)
