## Supplementary material for "Heterochromatin-dependent transcription of satellite DNAs in the *Drosophila melanogaster* female germline": TableS4

TableS4. *Rsp* expression level correlates with its copy number in the genome

| Method I: northern blot and slot blot | | |
| --- | --- | --- |
| strains | *Rsp* DNA repeat number estimated from Slot blot | *Rsp* RNA expression level quantified from Northern blot |
| ZW144 | 200 | 0.966552 |
| Ral357 | 600 | 6.798309 |
| Iso1 | 1100 | 7.934042 |
| Ral380 | 2300 | 8.456499 |
| It pk cn bw | 4100 | 17.28229 |
| Method II: qPCR and qRT-PCR | | |
| strains | *Rsp* DNA repeat number estimated from qPCR | *Rsp* RNA expression level quantified from qRT-PCR |
| ZW144 | 50 | 0.000195628 |
| Ral357 | 400 | 0.00190355633333333 |
| Iso1 | 1100 | 0.0050540994 |
| Ral380 | 2600 | 0.0061967918 |
| It pk cn bw | 5200 | 0.01619652775 |

Method I (Pearson’s correlation coefficient r^2^=0.93, p-value=0.02).

Method II (Pearson’s correlation coefficient r^2^=0.98, p-value=0.003).
